# Mechanical Checkpoint for Cell Division in Three-Dimensional Microenvironments

**DOI:** 10.64898/2026.06.16.732593

**Authors:** Md Foysal Rabbi, Donghyun Yim, Matthew Boyd, Sungmin Nam, Ovijit Chaudhuri, Taeyoon Kim

## Abstract

Cell division within mechanically confining extracellular matrices (ECMs) is a key regulator of tissue morphogenesis and cancer progression. Although the intracellular force-generation mechanisms that drive volumetric growth and mitotic elongation are well characterized, how ECMs resist these forces remains poorly understood. Unlike linearly elastic materials, fibrillar ECMs exhibit nonlinear and viscoelastic behaviors that fundamentally alter how they oppose cell-generated stresses. Using a fiber-level computational model, we dissected the origins of ECM-mediated mechanical confinement during mitosis. We identified three distinct modes of resistance: compressive resistance at the cell poles, shear resistance from a pericellular shell, and tensile resistance at the cell equator. The relative contributions of these modes depended on fiber architecture and connectivity; however, shear resistance from the pericellular shell—pre-tensed by volumetric growth during G1—consistently dominated as the primary mechanical barrier to mitotic elongation. These findings suggest that the pericellular shell functions as a natural mechanical checkpoint on cell division within collagen-rich microenvironments. Notably, a finite element continuum model, despite being the most widely used framework for tissue mechanics, failed to reproduce these behaviors, underscoring the necessity of fiber-resolution approaches. We propose that overcoming this mechanical checkpoint is a critical step in cancer progression, enabling cells to divide within the dense stromal matrices characteristic of metastatic tumors.

## INTRODUCTION

Cell division is a highly coordinated and essential biological process, mediating tissue development, regeneration, and homeostasis across multicellular organisms (*1*). Successful division requires not only accurate chromosome segregation but also substantial changes in cell shape. Cells typically double their volume prior to undergoing division, with much of the growth occurring during the G1 phase (*2, 3*), followed by mitotic rounding, a well-studied process, prior to division (*4, 5*). Then, cells undergo elongation along the mitotic axis, or mitotic elongation, a process critical for proper segregation of chromosomes and other intracellular materials and for guiding orientation of daughter cells (*6, 7*). This mitotic elongation is driven by intracellular mechanisms including interpolar spindle elongation (ISE) involved with the outward protrusion of cell poles (*8–12*) and cytokinetic ring contraction (CRC) that constricts the cell equator (*13–16*). These mechanisms have been well-characterized by isolated cell systems using flat two-dimensional (2D) substrates or in suspended cultures.

However, these traditional experimental protocols do not accurately account for the complex, confining environments that dividing cells experience in vivo. In tissues, cells are often embedded in dense extracellular matrices (ECMs) composed primarily of fibrous proteins such as type I collagen (*17, 18*). These matrices exert mechanical resistance to changes in cell morphology. It was unclear whether the mechanisms of mitotic elongation observed in unconfined or weakly resistant environments still operate effectively with physiological constraints. To address this knowledge gap, we recently explored mitotic elongation occurring in three-dimensional (3D) collagen gels that mimic stromal environments (*11, 19*). By combining live-cell imaging with perturbations, we showed that outward pushing forces generated by ISE and CRC in dividing cells actively deform the surrounding matrix fibers to achieve the mitotic elongation (*11*). In addition, using a computational model, we demonstrated that both ISE and CRC are necessary to accomplish the mitotic elongation in dense ECMs with high stiffness. These results showed that the mitotic elongation in 3D is not merely a byproduct of internal cellular rearrangements but generates space in confining microenvironments for proper cell division.

However, understanding how ECM resists this space-generating process is complicated in nature due to its sophisticated mechanical properties. Unlike linear elastic materials often assumed in simplified theoretical and computational models, type I collagen networks exhibit highly nonlinear behaviors; they exhibit strain-stiffening, characterized by an increase in stiffness at large shear/tensile strain for inhibiting excessive deformation, and collagen fibers can be buckled under compression, leading to asymmetric mechanical responses (*20–22*). Furthermore, the transient nature of cross-linking points between collagen fibers results in viscoelasticity and plasticity, allowing the matrix to relax stress and irreversibly remodel over time (*23, 24*). These complicated properties suggest that mechanical confinement which a dividing cell encounters is spatially varying and dynamic.

In this study, we extend our work by delving into how ECM resists mitosis. We adopted a fiber-level matrix model from our previous work and incorporated cell growth, prior to mitotic elongation, which is characterized by a two-fold increase in cell volume. Due to its discrete nature, the model can naturally capture the key mechanical features of ECMs, such as strain stiffening and fiber buckling. Our model can also represent the viscoelasticity and plasticity of ECMs via the force-dependent breakage of cross-linking points between fibers. We simulated cell growth followed by mitotic elongation under various conditions of ECMs to identify how specific matrix properties contribute to resisting mitotic elongation.

## METHODS

### Brownian dynamics with the Langevin equation

In the agent-based model, elements are defined by nodes. The velocities of the nodes of all elements, d**r***i* /d*t*, are calculated by the Langevin equation with inertial effects neglected:

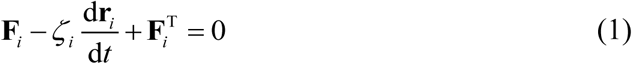

where **r***i* is the location of the *i*-th node, *ζi* is a drag coefficient, **F***i* is a deterministic force, and **F** ^T^ is a stochastic force satisfying the fluctuation–dissipation theorem (*25*):

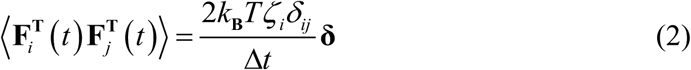

where *δij* is the Kronecker delta, Δ*t* = 3.97 ×10^-4^ s is time step, *k*B*T* is thermal energy, and **δ** is a unit second-order tensor. At each time step, the positions of the nodes of all elements are updated using the forward Euler integration scheme and velocities (d**r***i* / d*t*) calculated from Eq. 1:

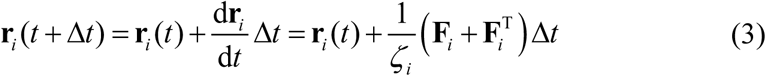

**F***i* includes extensional, bending, and repulsive forces. The extensional and bending forces are derived from the following harmonic potential functions:

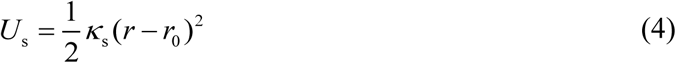

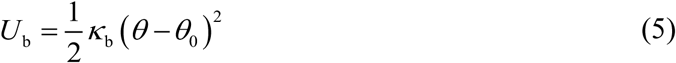

where *κ*s and *κ*b denote extensional and bending stiffnesses, respectively. *r* and *r*0 refer to instantaneous and equilibrium lengths, and *θ* and *θ*0 indicate instantaneous and equilibrium angles.

### Simplification, mechanics, and dynamics of fibers and cross-linkers

ECM is modeled as a matrix consisting of inter-connected fibers (Fig. 2A). Each matrix fiber is simplified into serially connected cylindrical elements. The equilibrium length of fiber elements (*r*0,f = 1 μm) and an equilibrium angle formed by adjacent fiber elements (*θ*0,f = 0 rad) are governed by extensional (*κ*s,f) and bending (*κ*b,f) stiffnesses of the fibers, respectively. The reference value of *κ*b,f corresponds to the persistence length of ∼100 μm.

Cross-linkers that physically connect matrix fibers comprise two cylindrical elements connected at their center point. The equilibrium length of cross-linker elements (*r*0,xl = 200 nm) is maintained by extensional stiffness (*κ*s,xl). An equilibrium angle between two arms of each cross-linker (*θ*0,xl,1 = 0 rad) and an equilibrium angle between a cross-linker arm and the axis of a fiber where the arm is bound (*θ*0,xl,2 = π/2 rad) are regulated by two bending stiffnesses (*κ*b,xl,1 and *κ*b,xl,2). Forces exerted on fiber elements by cross-linkers are distributed onto the two nodes of each fiber element. As the binding point is closer to a node, a larger portion of the forces is distributed to the node as described in our previous work (*26*).

*ζi* of fiber and cross-linker elements is estimated using an approximate formula for cylindrical objects (*27*):

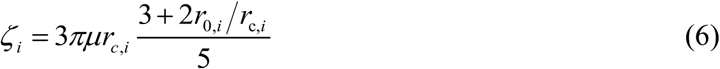

where *µ* is the viscosity of a surrounding medium, and *r*c,*i* and *r*0,*i* represent the diameter and length of elements, respectively.

At the beginning of simulations, there are a specified number of fiber elements in a monomer state. These monomeric fiber elements do not have any physical position; their local concentration is considered instead. The formation of fibers begins with the emergence of a single fiber element with the rate constant of *k*n,f. This element is extended by sequentially adding identical cylindrical segments in both directions with the rate constant of *k*+,f. The average length of the resulting fibers under the reference condition is ∼10 µm. Once fibers are assembled, they are not disassembled till the end of simulations. Like fiber elements, all cross-linkers are initially in a monomeric state at the beginning of simulations without any physical position. The cross-linkers bind to binding sites located every 100 nm on each fiber element with the rate constant of *k*+,xl. Then, these cross-linkers have their position and can bind to other neighboring fiber with the same rate constant. They can also unbind from fibers with a force-dependent rate, *k*-,xl, determined by Bell’s law [3]:

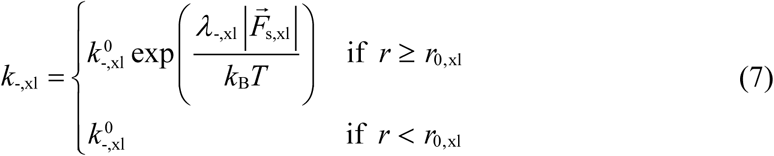

where *k_-,x1_*^0^ denotes the zero-force unbinding rate constant, *λ*-,xl indicates sensitivity to an applied force, *k*B*T* represents thermal energy, and 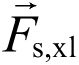 is a spring force vector acting on the cross-linker element. Only when the spring force is a tensile force, *k*-,xl becomes higher than its base rate, *k_-xl_*^0^. After each unbinding event, cross-linkers remain bound on one fiber and wait for binding to other fibers.

### Simplification and mechanics of the cell membrane

A cell membrane, which separates an intracellular space from an extracellular space, is simplified into a 2D or 3D structure, depending on the dimension of simulations. The 2D cell membrane is composed of interconnected rectangular elements. The nodes of the rectangular elements have only x and y positions; the rectangular elements span across the entire domain in z direction. The equilibrium length of the membrane elements (*r*0,m = 400 nm) is maintained by extensional stiffness (*κ*s,m). An equilibrium angle between adjacent membrane elements (*θ*0,m = 0 rad) is maintained by bending stiffness (*κ*b,m).

Repulsive forces between the membrane elements and elements representing fibers and cross-linkers keep the matrix at the outside of the membrane and also result in matrix deformation when the membrane changes its shape. A minimum distance between a membrane element and a cylindrical segment representing fibers or cross-linkers, *r*12, is computed. Then, a repulsive force is determined by the following potential:

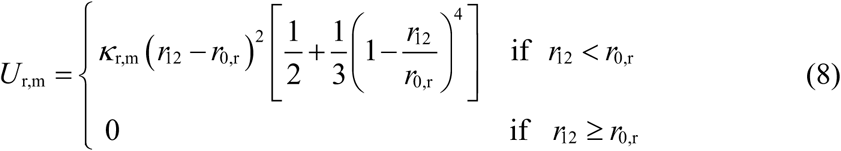

where *κ*r,m is the strength of the repulsive force, and *r*0,r is a distance below which the repulsive force starts acting. In this case, *r*0,r is equal to the average thickness of the membrane element (*r*c,m) and the cylindrical element. Note that this repulsive force increases more rapidly as *r*12 approaches *r*0,r. The repulsive force also acts between neighboring membrane elements, which is determined by Eq. 8 with *r*0,r = *r*c,m.

Volume encapsulated by the membrane is conserved. The 2D membrane approximates the cross-section of a membrane undergoing cytokinesis. In simulations, the membrane shape remains nearly symmetric along its long axis during cytokinesis. Thus, with the axisymmetric assumption, the volume of a 3D cell is estimated by rotating the 2D membrane shape about the long axis. If instantaneous cell volume (*V*m) is not equal to initial volume (*V*0,m), forces are applied to all membrane elements inwards or outwards in normal directions to maintain *V*m close to *V*0,m, which originate from the following harmonic potential:

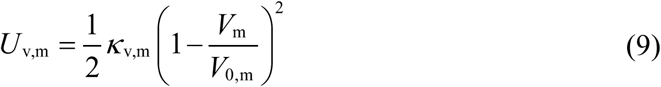

where *κ*_v,m_ is the strength of volume conservation.

The 3D cell membrane is modeled using a triangulated mesh. The equilibrium length of chains between mesh nodes (*r*0,m = 400 nm) is maintained by the same extensional stiffness (*κ*s,m). An equilibrium dihedral angle formed by adjacent triangular mesh elements (*θ*0,m = 0 rad) is regulated by the same bending stiffness (*κ*b,m). Volume encapsulated by the 3D membrane is conserved using Eq. 9. Repulsive forces act between neighboring triangular mesh elements and between triangular mesh elements and elements representing fibers and cross-linkers, using Eq. 8. *r*0,r is calculated in a similar manner, using the thickness of the triangular mesh elements (*r*c,m).

### Cell growth and mitotic elongation

To study mitotic elongation occurring within confining ECM, simulations are performed using either 2D or 3D setup. In the 2D setup, a thin cylindrical computational domain with 50 µm in radius and 1 µm in height (in the z direction) is employed. At the beginning of each simulation, a circular membrane with 10 μm in radius, which consists of 157 elements, is placed at the center of the domain. In the 3D setup, a spherical domain with 30 µm in radius is used, and a spherical membrane with 10 μm in radius, which consists of 10,242 nodes, is located at the domain center. A matrix is created between the membrane and the boundary of the domain via the self-assembly process of fibers and cross-linkers explained earlier. In the 2D setup, fibers are initially formed in the random orientation perpendicular to the z direction to prevent their elongation from being hindered by the small thickness of the domain. Cross-linkers bind to pairs of fibers to form cross-linking points. Note that the nodes of fibers, crosslinkers, and 3D membrane are free to move in the x, y, and z directions, whereas 2D membrane nodes move only in x and y directions because they do not have z positions.

After the matrix assembly phase for 100 s, cell growth begins by linearly increasing the equilibrium volume of the membrane two-fold for 4,000 s. Then, volume conservation described earlier (Eq. 9) pushes the membrane outwards uniformly to maintain volume within the membrane close to the increasing equilibrium volume (Fig. 2B). To increase the two-fold volume increase, the radius of the membrane increases up to ∼1.26-fold. After the cell growth phase, mitotic elongation starts by applying two types of forces to the membrane as done in our previous study (Fig. 2C) (*11*); to replicate contractile forces exerted on the cell equator by the CRC, a constant inward force (*F*CRC) is uniformly applied to membrane nodes within 0.5 µm from the equator. As a result, the membrane nodes subjected to this force move inward until their distance to the mitotic axis is greater than 1 µm. To account for the expansion of the ISE, an outward force (*F*ISE) is uniformly applied to membrane nodes within 5 µm from the mitotic axis. These two forces stop being applied if a cell elongated by 58.74 % to prevent two daughter cells from elongating more than a spherical shape.

### Finite element modeling

Cell growth and mitotic elongation are also simulated by finite element (FE) simulations (ANSYS Academic Research Mechanical, release 22.2) to directly compare continuum-level predictions with those from fiber-level ones. All geometric dimensions are chosen to be consistent with those used in the fiber-level model (Fig. 8A). The computational domain represents a quarter of the system composed of a cell surrounded by a matrix. A circular region at the center of the domain with 10 µm in radius represents a cell. An annular region with 50 µm in radius, which surrounds the circular region, represents ECM. In some of the simulations, a thin annular region with 10.5 µm in radius is added between the cell and the ECM as a shell (Fig. 8E). No overlap exists between these regions. All regions have identical thickness, 1 µm, and symmetric boundary conditions are applied along the x- and y-axes for symmetric conditions. The interface between the cell and its adjacent region (the thin layer or the matrix) is assumed to be frictionless, whereas the interface between the shell and the matrix is assumed to be bonded. Both the cell and the matrix obey the Neo-Hookean hyperelastic constitutive laws. The cell is treated as an incompressible solid with the initial elastic modulus of 324 Pa (*28*). The ECM is treated as a compressible solid with material parameters derived from type I collagen gels (*29*); its elastic modulus (*E*mat) is 832 Pa, and its mechanical compressibility is defined by either the Poisson’s ratio of 0.313 or equivalently the bulk modulus of 741 Pa. The elastic modulus of the shell (*E*shell) is equal to *E*mat or 10-fold higher than *E*mat. To mimic cell volumetric growth, constant surface tractions are applied to the cell boundary. To account for the CRC, inward forces are applied along the equatorial region of the cell, whereas outward forces are applied to the polar regions to mimic the ISE. These are consistent with loading configurations used in the fiber-level simulations (Figs. 1B, C).

**Figure 1.**
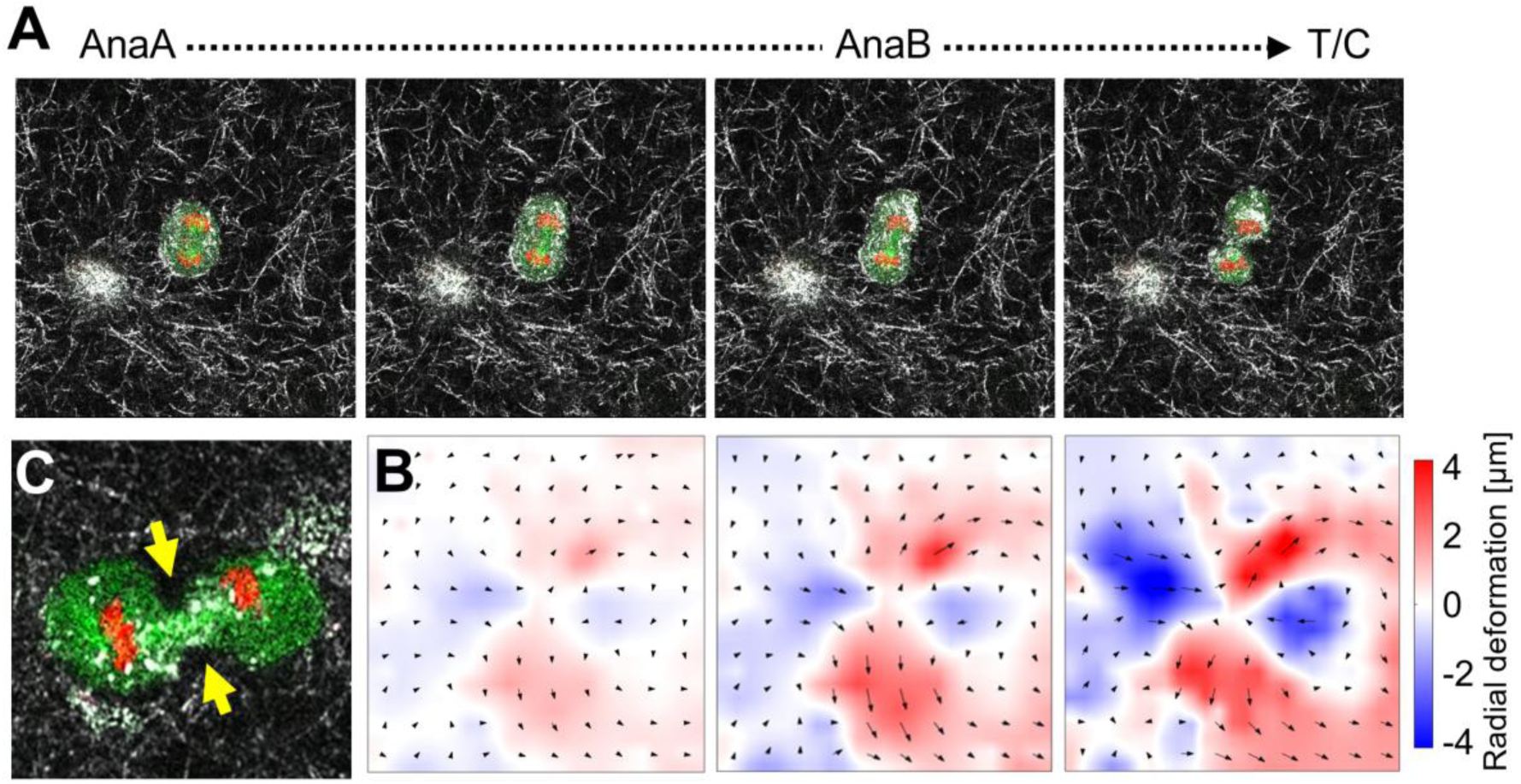
Mitotic elongation generated pericellular voids and anisotropic deformation in three-dimensional (3D) collagen matrices. (A) Time-lapse fluorescence images of an MDA-MB-231 cell dividing within a 3D collagen gel. Cells express GFP-α-tubulin (microtubules, green) and RFP-histone (chromatin, red), overlaid with collagen reflectance imaging (gray) to visualize the matrix. (B) Matrix deformation fields during division, illustrating outward deformation near the cell poles and inward deformation near the equator. This highlights spatially heterogeneous mechanical interactions between the dividing cell and the matrix. (C) At the late stage of mitotic elongation, cleavage furrow ingression and daughter-cell separation induced pericellular matrix voids (yellow arrows), indicating local detachment between the cell surface and the surrounding matrix.

**Figure 2.**
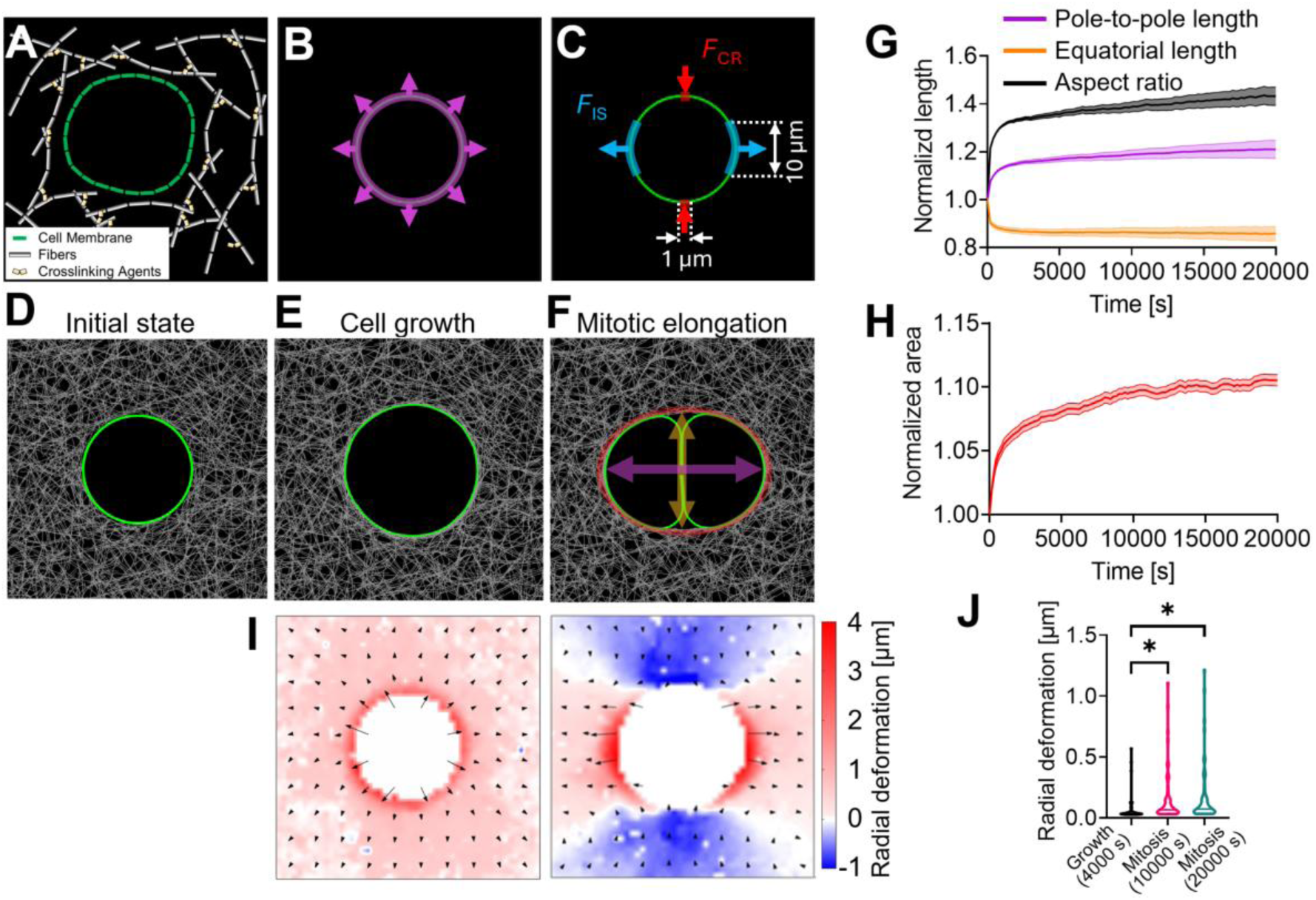
Discrete fiber matrix modeling recapitulated mitotic elongation and anisotropic matrix deformation. (A) Schematic of the computational framework. A matrix is represented as a discrete network consisting of serially connected fiber segments (gray) linked by dynamic cross-linkers (yellow), and the cell boundary is represented as a closed loop of interconnected membrane elements (green). (B) Pre-mitotic growth, simulated by applying uniformly outward expansile forces to the membrane to mimic isotropic volume increase during G1. (C) Mitotic elongation, simulated by applying an inward equatorial contractile force (exerted by the cytokinetic ring) and an outward force along the mitotic axis (exerted by interpolar spindle elongation). (D–F) Snapshots taken at three stages: initial state (D), cell growth (E), and mitotic elongation (F). (G) Time evolution of cell shape during mitotic elongation. Pole-to-pole length and equatorial length, indicated by horizontal and vertical arrows in (F) respectively, are normalized by their initial values. The aspect ratio represents the pole-to-pole length divided by the equatorial length. The cell rapidly elongates along the mitotic axis while narrowing at the equator, resulting in an increasing aspect ratio over time. (H) Temporal profile of a normalized area change in the pericellular shell during division. (I) Matrix deformation fields during growth and mitosis, reproducing the experimental outward deformation at the poles and inward deformation at the equator. Radial deformation was quantified by projecting local fiber displacement vectors onto the outward radial direction from the cell center. (J) Quantification of radial deformation at representative times during growth and mitosis. Data in (G, H) are presented as mean ± s.d. (*n* = 4 independent simulations). Statistical analysis in (J) was performed using the one-way ANOVA with multiple-comparisons correction across the groups shown in (J). *: p < 0.05.

### Experimental protocols

To validate our computational model, we utilized experimental datasets derived from previously published studies (*11*). The experimental platform employed MDA-MB-231 human breast cancer cells that are genetically modified to express GFP-labeled α-tubulin and RFP-labeled histone, enabling high-resolution imaging of spindle microtubules and chromatin, respectively. These cells are cultured in high-glucose Dulbecco’s Modified Eagle Medium (DMEM) supplemented with 10% fetal bovine serum and 1% penicillin/streptomycin. To probe cell division dynamics, cells are embedded in type I collagen gels with final concentrations of 1 mg/mL. The gels are prepared by neutralizing collagen stock solution (rat tail, pH adjusted to 8.5 using NaOH) in the presence of suspended cells at a density of 0.1–0.5 million cells/mL. Polymerization is carried out at 37 °C for 45–60 min in glass-bottom plates suitable for high-resolution imaging.

Live-cell imaging is performed using a Leica SP8 confocal microscope under standard culture conditions (37 °C, 5% CO₂). For time-lapse imaging of mitotic progression, cells are monitored using a 25×/0.95 NA water-immersion objective. Fluorescence signals are acquired using excitation at 488 nm for GFP-labeled microtubules (α-tubulin), 555 nm for RFP-labeled histone, and 639 nm for collagen fiber reflectance. Quantitative image analysis is performed using ImageJ. Matrix deformation fields are computed by tracking collagen reflectance fibers using a particle image velocimetry (PIV) algorithm (PIVlab in MATLAB). Erroneous or unstable displacement vectors, especially those corresponding to regions inside the cell body, are corrected by averaging nearby vectors.

### Statistical Analysis

For each condition, we ran 4 independent simulations, unless otherwise specified. In some of the plots, data are presented as mean ± standard deviation (s.d.). For comparisons between more than two groups, we performed one-way analysis of variance (ANOVA) followed by Tukey’s multiple-comparison test. For pairwise comparisons, unpaired two-tailed Student’s t-tests were used. Statistical significance is indicated as follows: ns: p ≥ 0.05, *: p < 0.05, **: p < 0.01, ***: p < 0.001, ****: p < 0.0001.

## RESULTS

### Mitotic elongation induced anisotropic matrix deformation in 3D collagen networks

First, we performed 3D culture experiments using MDA-MB-231 cells embedded within type I collagen gels and observed how cell morphology changed during mitosis. As the cells progressed from anaphase A (AnaA) into telophase/cytokinesis (T/C), they underwent pronounced elongation along the mitotic axis. During this mitotic elongation phase, the aspect ratio of the cell body increased significantly (Fig. 1A). This anisotropic shape change induced spatially distinct matrix deformation, with outward displacement of collagen near the cell poles and inward deformation near the equator (Fig. 1B). Toward the end of mitotic elongation, as the cleavage furrow formed for separation into the two daughter cells, we observed the emergence of pericellular voids in the matrix near the equatorial plane (Fig. 1C).

### Model overview

To dissect the origin of mechanical confinement to mitosis, we employ a discrete model consisting of a cell dividing within a fibrous matrix, which is based on Brownian dynamics with the Langevin equation (Fig. 2A). Unlike continuum-based models, this modeling framework treats ECM as a matrix composed of interconnected semiflexible fibers. Each fiber is discretized into serially connected cylindrical elements experiencing extensional and bending forces. This discretization enables the matrix to capture nonlinear responses including strain-stiffening caused by tension/shear and fiber buckling induced by compression. Fiber connectivity and viscoelasticity are regulated by dynamic cross-linkers that reversibly connect pairs of fibers. The force-dependence of their dissociation rate enables the matrix to relax forces and allow fibers to be reoriented to a larger extent. A cell undergoing mitosis is described by a 2D or 3D membrane with 10 µm in radius. In a large number of 2D simulations, rectangular elements form a circular wall as the 2D membrane in a thin cylindrical domain (50 µm in radius and 1 µm in height). In a single 3D simulation which was performed to verify our results from the 2D simulations, a spherical membrane is described by a triangulated mesh in a spherical domain (30 µm in radius). Note that the computational cost of this 3D simulation is ∼ 6-fold higher than that of the 2D simulations; the total number of elements in the 3D simulation is 1,093,992, whereas that in the 2D simulations is only 180,474.

To mimic forces generated by dividing cells, we considered two-fold volumetric growth, driven by isotropic internal pressure, and mitotic elongation, driven by expansile forces from elongating interpolar spindle and localized contractile forces from the cytokinetic ring constricting at the cell equator (Figs. 2B, C).

### Discrete modeling recapitulated mitotic elongation observed in experiments

To consider the effect of physical memory in a matrix on confinement to mitotic elongation, we simulated both cell growth and mitotic elongation occurring in a fibrillar matrix. We first emulated cell growth by applying uniform outward forces to the membrane via volume conservation (Fig. 2B). Then, two types of forces are applied to the two poles and equator of the cell to mimic mitotic elongation (Fig. 2C). As a result of the force application, the cell underwent two-fold volumetric expansion and then an increase in its aspect ratio (Figs. 2D-F). During the mitotic elongation, a cell body elongates by ∼20 % in one direction and shrinks by ∼15 % in the other direction, resulting in the aspect ratio of 1.41 (Fig. 2G). We also observed voids between the cell membrane and the matrix near the cell equator as in experiments (Fig. 2F). These voids emerged because a pericellular shell could not conform to the inward movement of the membrane. We found that this shell experienced a ∼10% increase in its effective surface area during mitotic elongation (Fig. 2H).

As the cell changes its shape, the surrounding matrix was also deformed. Volumetric growth deformed the surrounding matrix, but this deformation was highly localized near the cell membrane rather than propagating over long distances, creating a zone with high matrix density (Fig. 2I, left). This localization arose because many of the matrix fibers buckled in response to compressive loads beyond a buckling load. Consequently, this growth phase effectively formed a densified pericellular shell. Mitotic elongation also led to outward matrix deformation near cell poles and inward deformation near cell equator (Fig. 2I, right), which is similar to experimental observations (Fig. 1B). It was found that matrix displacement was higher during the mitotic phase compared to the growth phase (Fig. 2J). Overall, the significant matrix deformation indicates that a dividing cell conducts active mechanical work against the ECM to generate a sufficient space required for elongation.

### A matrix resists mitotic elongation through three distinct mechanical modes

To identify the physical mechanism by which the ECM opposes cell division, we analyzed the spatiotemporal evolution of forces developed on the matrix. At the end of the growth phase, outward cell expansion generated a concentrated buildup of tensile forces near the cell boundary, forming a shell-like structure under high tension (Fig. 3A). Tensile forces along the shell were nearly uniform (Fig. 3B). This isotropic yet prestressed state serves as an initial mechanical condition from which spatially distinct resistance modes emerge during mitotic elongation. As the cell start elongating, high compressive forces developed near the poles because these regions of the matrix were directly pushed by the elongating cell (Fig. 3D, right). Concurrently, tensile forces were concentrated on a subset of matrix fibers forming a pericellular shell that wraps around the cell (Fig. 3D, left). These tensile forces, together with the simultaneous areal expansion of the shell, demonstrate that it functions as a pre-tensioned hoop-stress cage that acts as a primary mechanical checkpoint against the geometric changes required for mitosis. Interestingly, tensile forces measured along the shell were not uniform; it peaked at the two poles and diminished near the equator (Fig. 3E), meaning spatial anisotropy in mechanical resistance from the shell. This anisotropy arises from physical connections between the shell and its surroundings; a small number of matrix fibers near the equator bore disproportionately high tensile loads because they serve as anchoring elements that couple the shell to the rest of the matrix. These anchors resist changes in the aspect ratio of the shell during mitotic elongation.

**Figure 3.**
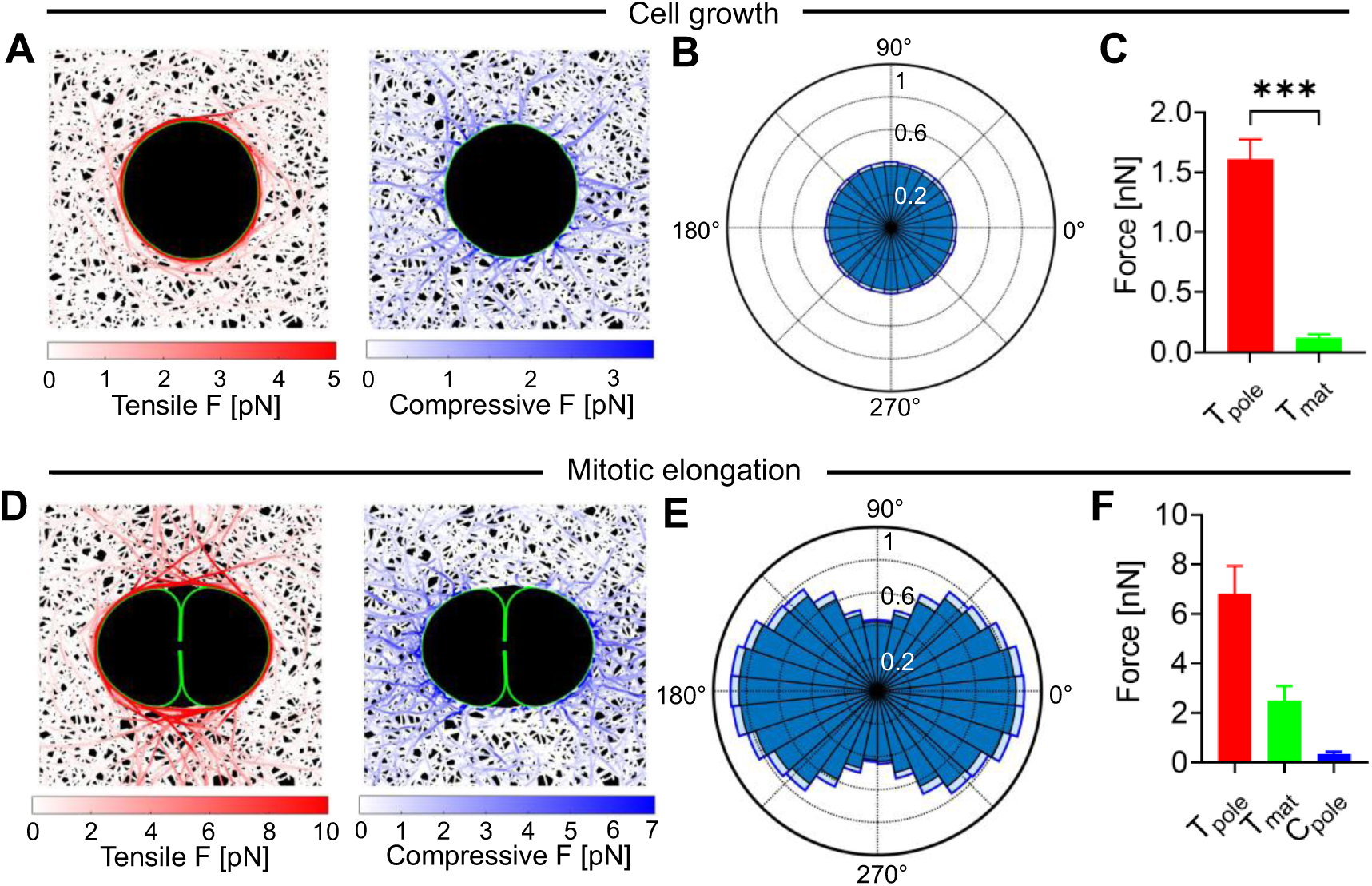
The extracellular matrix resists mitotic elongation through growth-induced shell tension, polar compression, and matrix anchoring. (A) Snapshots showing the spatial distribution of tensile and compressive matrix forces at the end of cell growth. Tensile forces were concentrated along the pericellular shell, whereas compressive forces were distributed around the cell boundary. (B) Angular distribution of tensile forces along the shell at the end of cell growth, demonstrating a nearly uniform distribution. (C) Quantitative comparison of tensile resistance components at the end of cell growth: *T*pole denotes the tensile force measured on the shell near the poles, whereas *T*mat represents the relative contribution of anchoring between the shell and a bulk matrix. (D) Snapshots showing tensile and compressive forces during mitotic elongation. Tensile forces were concentrated along the pericellular shell and anchoring fibers near the cell equator, whereas compressive forces were prominent near the poles. (E) Angular distribution of tensile forces measured along the shell during mitotic elongation, showing peak tension near the poles that decreased toward the equatorial region. (F) Decomposition of matrix resistance into three components: *T*pole, *T*mat, and compressive forces near the poles (*C*pole). *T*mat is equal to a difference between *T*pole and the tensile force measured on the shell near the cell equator. *T*mat was ∼37% of *T*pole, implying the important role of the anchoring in tension distribution. Data in (B, C, E, F) are presented as mean ± s.d. (*n* = 4 independent simulations). Statistical analysis in (C) was performed using two-sided unpaired t-tests (C). ***: p < 0.001.

In sum, we identified three sources of matrix resistance to mitotic elongation: compressive resistance near the cell poles, resistance to areal expansion of the pericellular shell, and tensile resistance opposing the inward deformation of the shell. We quantified compressive forces acting near the two poles (*C*pole) and tensile forces exerted on the shell near the poles (*T*pole) (Fig. 3F). We also measured tensile forces acting on the shell near the equator (*T*eqt). If the shell were mechanically isolated from the surrounding matrix, *T*pole would be comparable to *T*eqt due to force balance. Therefore, a difference between them, *T*mat = *T*pole - *T*eqt, indicates the contribution of resistance transmitted through the connections between the shell and the rest of the matrix. In the reference case, *T*mat was ∼37% of *T*pole. Note that *T*pole and *T*eqt were similar to each other at the end of cell growth (Fig. 3C).

We ran additional simulations with the pericellular shell mechanically isolated; all fibers within an annular region whose thickness is 3 µm are severed to separate the shell from the bulk matrix (Figs. S1A-C). The isolated shell developed uniform tensile forces during both growth and mitotic elongation (Fig. S1D), which is contrast to anisotropic force development observed on the shell connected to the bulk matrix (Fig. 3E). *T*pole was comparable to *T*eqt as can be inferred from negligible *T*mat (Fig. S1E), confirming that the shell behaved as an isolated shell. Cell elongation was noticeably enhanced when the shell was isolated, which could be attributed to less compressive resistance at the poles (Fig. S1C) and the absence of anchors to the bulk matrix. It is likely that the latter was a more significant cause for the enhanced cell elongation. This implies that physical connections between the shell and the bulk matrix play an important role in ECM resistance to mitotic elongation.

In addition, to evaluate the role of pre-mitotic cell growth in establishing matrix resistance, we performed simulations in which mitotic elongation occurred without a preceding growth phase. In the absence of growth, both the *T*pole and *T*mat were markedly reduced (Figs. S2A, B). This drastic reduction confirms that the pericellular shell is not merely a passive structural feature but a stress-hardened barrier generated specifically by the volumetric expansion of the G1 phase. Thus, the growth phase functions as a critical mechanical checkpoint, transforming the ECM into a high-resistance shell capable of bearing substantial tensile loads during mitosis.

### Tensile straightening and compressive buckling define spatially distinct matrix responses

We further investigated how fibers respond to mechanical stress. In live-cell experiments, we observed two different types of fiber deformation: fibers proximal to the cleavage furrow straightened during cytokinesis, whereas fibers near the poles increased their curvature, indicating that a substantial fraction of compressive forces were relaxed via buckling during mitotic elongation (Figs. 4A, B). Our simulations reproduced these structural phenotypes. Significant fiber deformation was observed only near the peripheral of the pericellular shell; proximal polar fibers exhibited a sharp increase in curvature consistent with buckling (Fig. 4C), whereas proximal equatorial fibers showed a pronounced reduction in curvature consistent with straightening (Fig. 4D). By contrast, distal fibers in both regions remained unchanged, confirming that matrix deformation induced by the dividing cell was highly localized.

**Figure 4.**
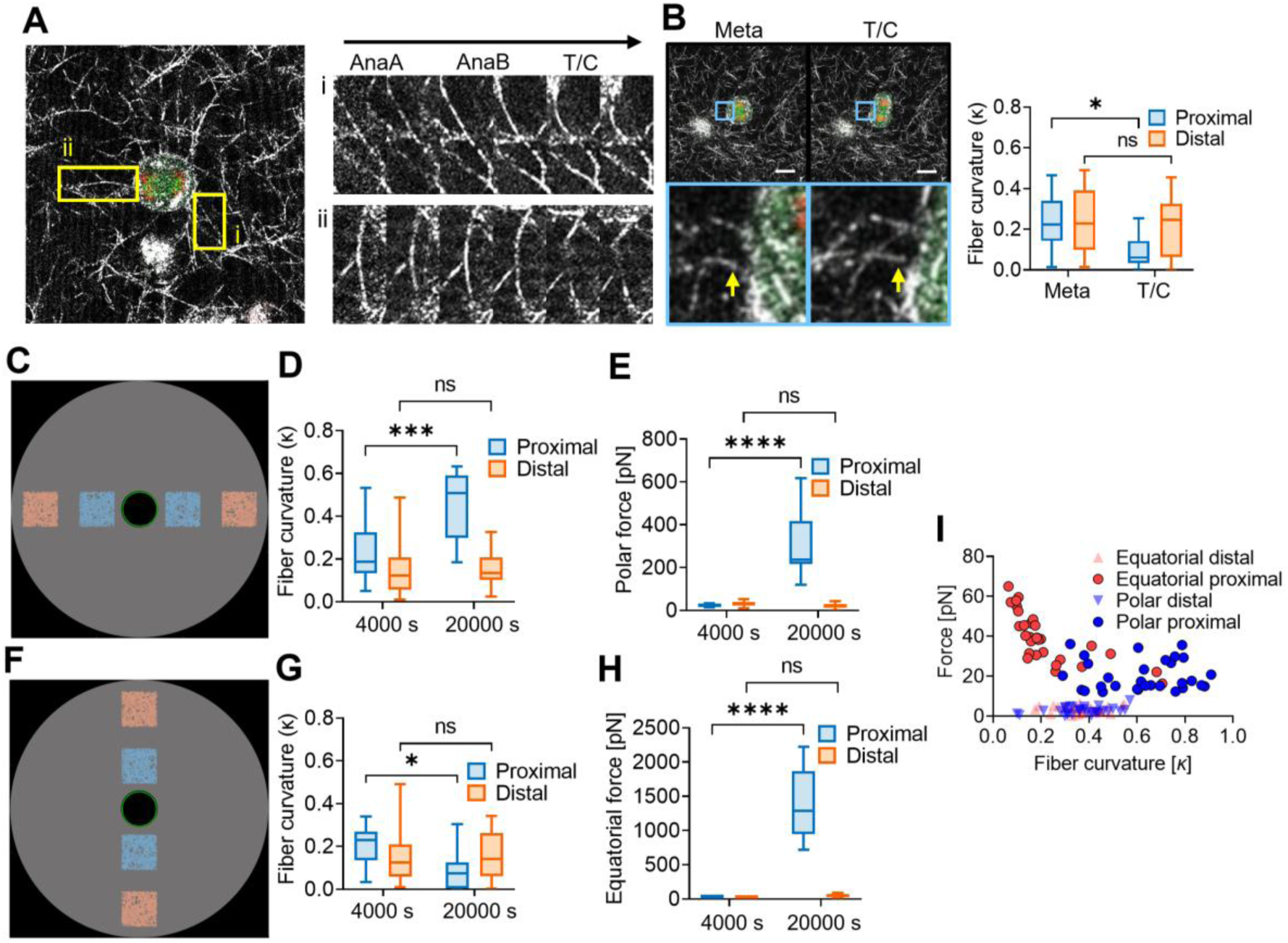
Spatially distinct matrix mechanics during mitotic elongation: equatorial tensile straightening versus polar compressive buckling. (A, B) Experimental images tracking collagen fiber remodeling during mitotic progression. (A) Two yellow boxes on the left indicate representative buckled fibers near cell poles shown as time-lapse images on the right from anaphase A (AnaA) to telophase/cytokinesis (T/C). (B) Blue boxes indicate straightened fibers near the equatorial region during T/C after metaphase. A box plot on the right shows a significant decrease in the curvature of fibers (i.e., straightening) proximal to the cell membrane. (C-H) Analysis of the curvature and force of fibers in simulations at the end of cell growth (4,000 s) and during mitotic elongation (20,000 s). (C, D) Fibers proximal to the cell poles in two blue regions in (C) showed a moderate increase in their curvature after cell growth and then a significant increase during mitotic elongation unlike those in two red regions. (E) Sum of the force magnitudes measured in the 4 colored regions in (C). Fibers near the cell poles experienced an increase in their forces during mitotic elongation, which originated mostly from compressive resistance. (F, G) Fibers proximal to the cell equator in two blue regions in (F) showed a similar increase in their curvature at the end of cell growth, but the curvature dropped substantially during mitotic elongation, implying straightening. (H) Fibers near the equator showed a large increase in their forces during mitotic elongation, attributed to anchoring between the shell and the bulk matrix. (I) Force magnitude and curvature of individual fibers during mitotic elongation. Equatorial proximal fibers tend to have low curvatures and high forces since they serve as load-bearing anchors, whereas polar proximal fibers tend to have higher curvatures and relatively low forces capped by their critical buckling load. All fibers in distal regions experienced negligible forces. In (B, D, E, G, H), data are presented as mean ± s.d. (*n* = 4 independent simulations), and statistical analysis was performed using one-way ANOVA. ns: p ≥ 0.05; *: p < 0.05; ***: p < 0.001; ****: p < 0.0001.

We quantified how these structural changes are correlated with mechanical resistance. We found that the magnitudes of forces acting on fibers in the regions proximal to the poles and the equator increased during mitotic elongation (Figs. 4E, F). We also found that some of the relatively straight fibers with lower curvature near the equator bore large tensile forces (Fig. 4G), which are the fibers forming anchorage between the pericellular shell and the bulk matrix. Fibers near the poles showed a much smaller increase in their force magnitudes because many of them underwent buckling due to compressive loads. All force magnitudes that the proximal polar fibers experienced were less than ∼ 30 pN which is close to the critical buckling load.

Together, these results demonstrate that the ECM undergoes highly localized, mode-specific structural remodeling in response to mitotic forces, and that these deformations directly encode the mechanical resistance experienced by the dividing cell. Tensile loading of straightened equatorial fibers establishes strong anchorage to the bulk matrix, whereas compressive loading at the poles is largely dissipated through fiber buckling, limiting the buildup of polar resistance. The tight correspondence between fiber curvature changes and force magnitudes highlights how the ECM fibrillar architecture partitions tensile and compressive loads during mitotic elongation, revealing a mechanically adaptive microenvironment that shapes the resistance landscape encountered by the cell.

### 3D simulations validate the intrinsic transition from isotropic to anisotropic resistance

To confirm the universality of these mechanical modes, we extended our framework to a fully 3D matrix to test whether the transition from isotropic resistance to anisotropic resistance is an inherent property of the matrix. The 3D simulations recapitulated the full sequence of cell division: initial state, isotropic volumetric growth, and mitotic elongation (Figs. 5A and S3A). During growth, the cell nearly doubled its volume, compacting nearby fibers and generating an isotropically prestressed pericellular shell. Upon entering mitotic elongation, the cell extended by ∼60%, accompanied by a ∼20% increase in shell area (Figs. 5B, C). This produced the outward deformation of the matrix at the poles and inward deformation near the equator, whereas cell growth resulted in isotropic outward matrix deformation (Figs. 5D and S3B). The characteristic signatures of polar compression, shell hoop stress, and equatorial anchoring were also observed as in 2D simulations (Figs. 5E, F). The 3D simulations confirmed that our 2D simulations are able to capture matrix deformation emerging during cell division, and that spatially distinct matrix resistance and force patterns are intrinsic to fibrous architectures.

**Figure 5.**
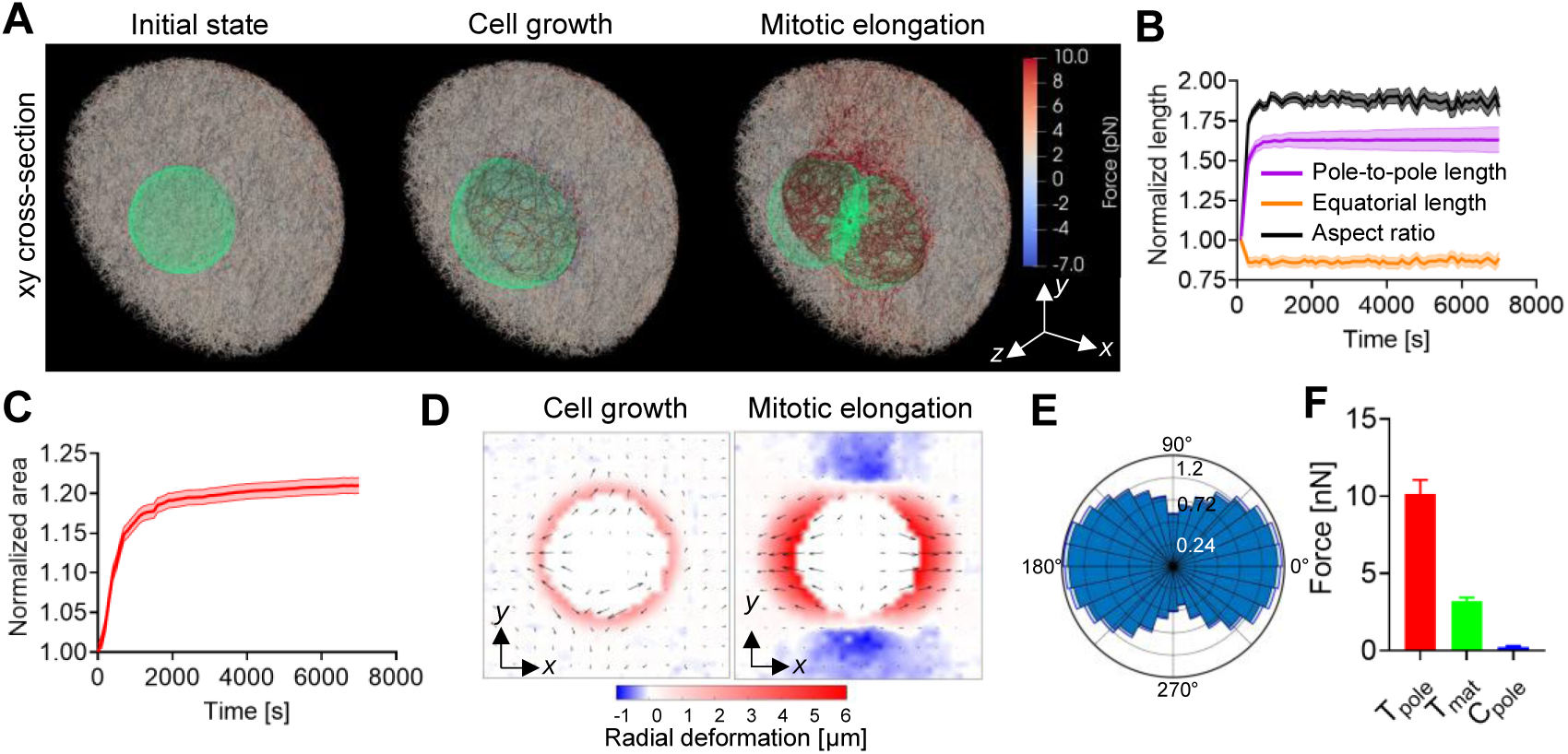
Three-dimensional (3D) simulations recapitulated growth-induced shell expansion, anisotropic matrix deformation, and force partitioning during mitotic elongation. (A) Snapshots showing three stages of cell division in 3D simulations: pre-growth, post-growth, and mitotic elongation. Images depict xy cross-sections. The cell boundary is shown in green, and color scaling on the right is used to visualize forces acting on matrix fibers. (B) Cell shape changes during mitotic elongation. (C) Time course of normalized pericellular shell area during mitotic elongation. (D) Matrix deformation fields at the end of the growth phase and during mitotic elongation. (E) Angular distribution of tensile forces along the pericellular shell during mitotic elongation. (F) Decomposition of matrix resistance into three components: shell tensile resistance (*T*pole), anchor resistance (*T*mat), and polar compressive resistance (*C*pole). All quantitative analyses were performed in the same manners as those for two-dimensional (2D) simulations. These results obtained in 3D simulations were qualitatively similar to those in 2D simulations although the 3D matrix was less confined than the 2D one shown in Figs. 2 and 3 due to different matrix conditions. Data in (B, C, E, F) are presented as mean ± s.d. (*n* = 4 independent simulations).

### Systematic tuning of matrix architecture reveals conditions for checkpoint engagement and failure

We systematically varied matrix properties to determine the conditions under which each mode of resistance to mitotic elongation becomes more or less dominant. For each parameter change, we quantified cell elongation, *T*pole, *T*mat, and *C*pole. Reducing fiber density (*C*f) from the reference case (Fig. 6A) weakened fiber-fiber connectivity and lowered overall matrix stiffness. As *C*f decreased, cell elongation increased (Fig. 6B), and all three resistive forces— *T*pole, *T*mat, and *C*pole—declined to a similar extent (Figs. 6C, 6D, S4A). This indicates that fiber density uniformly scales the global magnitude of the mechanical checkpoint without altering the balance between shell tension, polar compressive resistance, and equatorial anchoring. We next varied average fiber length (*L*f), another key determinant of network connectivity (Fig. 6E). Increasing *L*f enhanced both *T*pole and *T*mat (Figs. 6G and S4B). *C*pole increased in proportion to *L*f. Cell elongation was noticeably reduced, meaning that the cell was effectively locked into a highly confinement state (Fig. 6F). Conversely, halving *L*f markedly increased cell elongation and triggered a catastrophic failure of the mechanical checkpoint: *T*pole and *T*mat dropped sharply, whereas *C*pole decreased by less than half (Fig. 6G), revealing that tensile resistance is far more sensitive to fiber length than compressive resistance. The ratio *T*mat to *T*pole fell below 0.5 in short-fiber networks, indicating much weaker anchoring between the shell and the bulk matrix.

**Figure 6.**
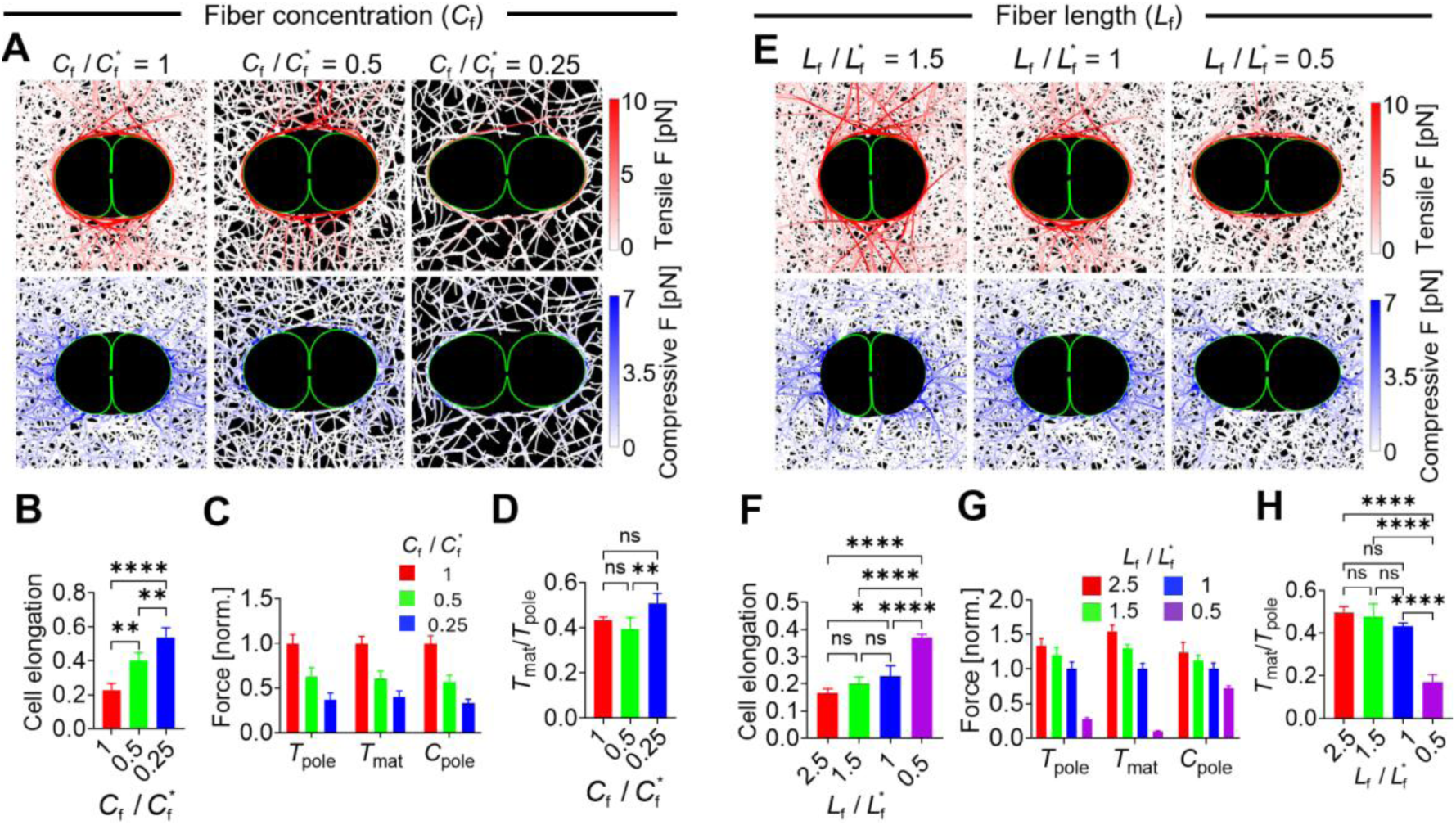
Lower fiber density enhanced cell elongation without a change in matrix resistance modes, whereas shorter fibers facilitated mitotic elongation by preferentially weakening pericellular anchoring. In (A-D), fiber density (*C*f) was reduced from its reference value ( *C*_f_*). In (E-H), fiber length (*L*f) was changed from its reference value ( *L*_f_*). (A, E) Snapshots showing the spatial distribution of tensile (top) and compressive (bottom) forces on the matrix. (B, F) Final cell elongation level. (C, G) shell tensile resistance (*T*pole), anchor resistance (*T*mat), and polar compressive resistance (*C*pole) normalized by their maximum values. (D, H) The ratio of *T*mat to *T*pole which represents the relative importance of anchors between the pericellular shell and the bulk matrix. Decreasing *C*f resulted in a sparser matrix network with reduced mechanical resistance and greater cell elongation. In addition, three types of resistance were reduced to similar extents. Lower *L*f also enhanced cell elongation, but the pericellular anchor between the shell and a bulk matrix played a much less important role when *L*f was reduced to half. Data in (B, C, D, F, G, H) are presented as mean ± s.d. (n = 4 independent simulations). Statistical significance in (B, D, F, H) was assessed using one-way ANOVA. ns, p ≥ 0.05; *, p < 0.05; **, p < 0.01; ****, p < 0.0001.

Cross-linking density (*R*xl) also modulates fiber connectivity. When *R*xl was reduced to half of the reference condition, cell elongation nearly doubled (Figs. S5A, B). Both *T*pole and *C*pole decreased substantially, whereas *T*mat showed only a modest reduction, leading to a slightly higher ratio of *T*mat to *T*pole. (Figs. S5C-E). It means that connection between the shell and the bulk matrix was less affected by the decreased cross-linking density. With further decreases in *R*xl, all three resistive forces— *T*pole, *T*mat, and *C*pole—declined, but *C*pole dropped more rapidly than the tensile components. This higher sensitivity of *C*pole to cross-linking density is expected, as increased cross-linking level reduces the effective buckling length by introducing more stabilized points along each fiber (*30*). We next tested if the mechanical checkpoint could be released by tuning the transient dynamics of cross-linkers. We increased one of the two parameters governing the force-dependent unbinding rate, *k*-,xl, up to 100-fold. Higher *k*-,xl increased cell elongation (Figs. 7A, B) and reduced *T*pole, *T*mat, and *C*pole to similar extents (Figs. 7C, S4C). However, the ratio *T*mat to *T*pole approached zero at the highest unbinding rate, indicating that the shell deformed with virtually no resistance from the bulk matrix (Fig. 7D).

**Figure 7.**
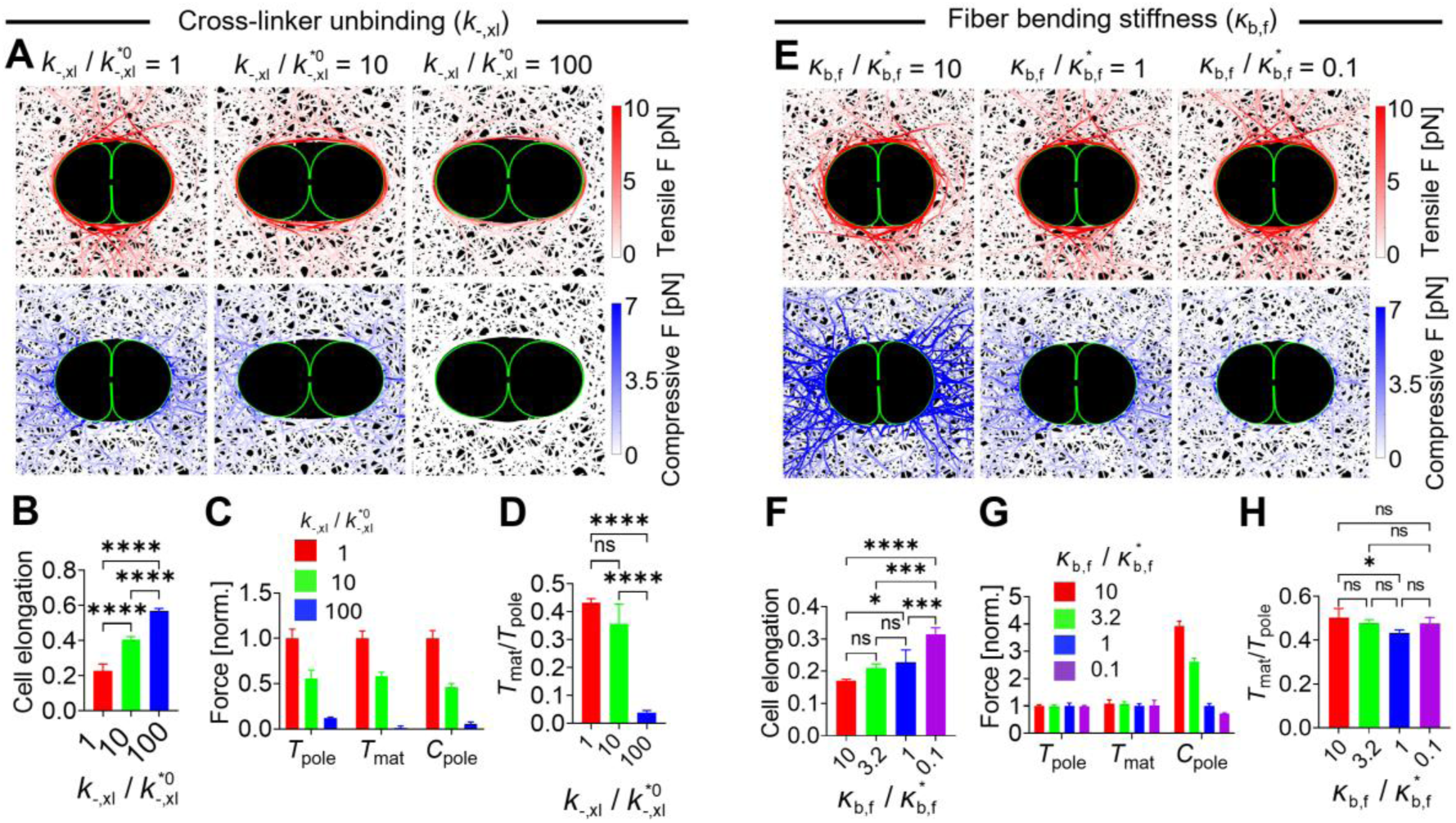
Frequent cross-linker dissociation from fibers promoted mitotic elongation by reducing overall matrix resistance, anchoring in particular, whereas increased fiber bending stiffness inhibited elongation by enhancing polar compressive resistance. In (A-D), the unbinding rate of cross-linkers 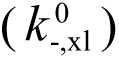 was increased from its reference value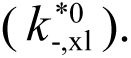 In (E-H), fiber bending stiffness (κ_b,f_) was varied from its reference value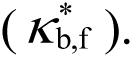 (A, E) Snapshots visualizing the spatial distribution of tensile (top) and compressive (bottom) forces on the matrix. (B, F) Cell elongation level measured in steady states. (C, G) shell tensile resistance (*T*pole), anchor resistance (*T*mat), and polar compressive resistance (*C*pole) normalized by their maximum values. (D, H) The ratio of *T*mat to *T*pole indicating the relative significance of anchors between the pericellular shell and the bulk matrix. Increasing 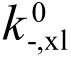 enhanced cell elongation by reducing all three types of mechanical resistance, but the pericellular anchors became much less important when 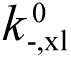 was very high. Increasing κ_b,f_ decreased cell elongation by enhancing only compressive resistance at cell poles. Data in (B, C, D, F, G, H) are presented as mean ± s.d. (n = 4 independent simulations). Statistical significance in (B, D, F, H) was assessed using one-way ANOVA. ns, p ≥ 0.05; *, p < 0.05; ***, p < 0.001; ****, p < 0.0001.

**Figure 8.**
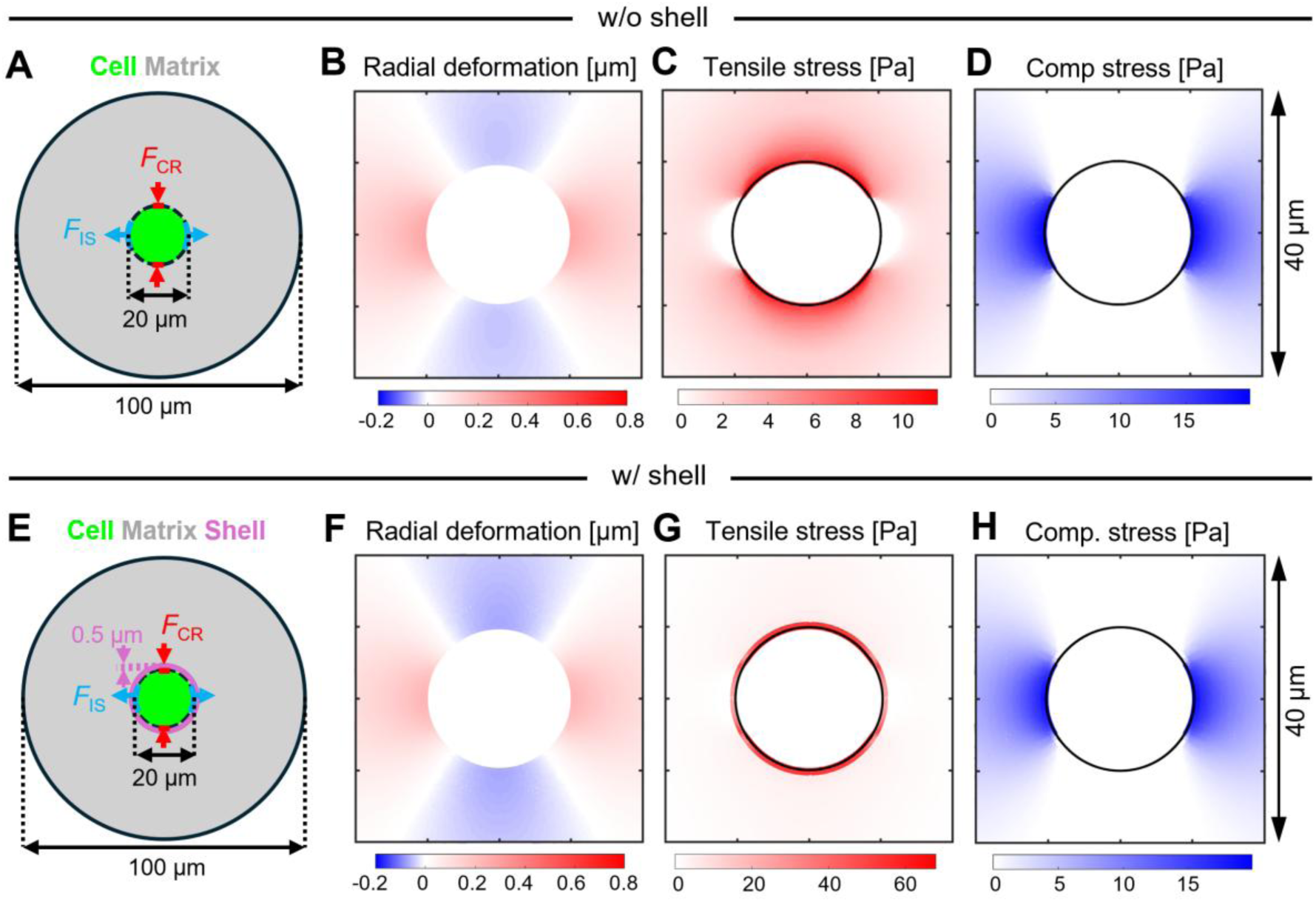
Finite element (FE) approaches reproduced global matrix deformation but failed to capture the distinct modes of matrix resistance. (A) Schematic of the FE model without a pericellular shell. The cell is directly surrounded by a matrix, and mitotic elongation is modeled by applying an inward contractile force at the equator (*F*CR) and an outward force along the mitotic axis (*F*IS). (B–D) Matrix radial deformation and stress distributions during mitotic elongation without a pericellular shell. Radial deformation and compressive stress distribution were similar to the patterns observed in discrete fiber-level simulations. However, tensile stress distribution failed to capture the pericellular shell. (E-H) The FE simulation was repeated in the presence of a thin pericellular shell with 0.5 μm in thickness and 10-fold higher stiffness than the surrounding matrix. Including the shell resulted in tensile stress distribution localized on the shell but failed to reproduce the coexistence of polar compressive resistance, shell resistance, and anchor resistance that emerged in discrete simulations. These results indicate that the three types of resistance are emergent microstructural properties not captured easily by continuum-based approaches.

We also probed the contribution of fiber stiffness. First, fiber bending stiffness (*κ*b,f) was varied over a wide range. Increasing *κ*b,f reduced cell elongation (Figs. 7E, F). While *T*mat and *T*pole were insensitive to *κ*b,f, *C*pole scaled in proportion to *κ*b,f (Figs. 7G, S4D). The critical buckling load is directly proportional to *κ*b,f. Thus, with higher *κ*b,f, enhanced compressive resistance at the poles suppresses cell elongation. We also reduced fiber extensional stiffness (*κ*s,f) from the reference condition. Lowering *κ*s,f to 1/25 of the reference value increased cell elongation and decreased *T*pole, *T*mat, and *C*pole to similar extents (Figs. S6A-D). However, with further reduction, *T*mat approached zero, indicating that the shell could deform without bulk-matrix resistance (Fig. S6E). Although *C*pole continued to decline, its rate of decrease was smaller. This seemingly counterintuitive dependence of compressive resistance on *κ*s,f arises because fibers are connected to other fibers at multiple points: when supporting fibers are easily extensible, stability is reduced, weakening the ability of fibers to resist compressive loads.

### Continuum-based approaches failed to reproduce multimodal resistance patterns in fibrous matrices

To determine whether these resistance modes can be captured by macroscopic continuum models, we performed finite element (FE) simulations using standard hyperelastic constitutive laws (Fig. 8A). Baseline FE simulations with spatially uniform material properties produced smooth, spatially continuous force and displacement fields around the dividing cell. The imposed equatorial contraction and polar protrusion resulted in outward radial matrix displacement at the poles and inward displacement near the equator (Figs. 8B, S7A)—consistent with the global deformation observed in the discrete simulations (Fig. 2I, right). However, the FE-generated force fields lacked the sharp spatial heterogeneity and directional contrasts observed in the discrete matrix (Figs. 3A, B and 8C, D); tensile forces decayed smoothly with distance from the cell, with no tension development at the poles (Fig. 8C). Circumferential tension and radial compression did not coexist within the same region, and no localized compressive relaxation zones analogous to fiber buckling were observed (Fig. S7A). Thus, while the FE baseline reproduced the qualitative global displacement pattern, it failed to capture the discrete mechanical signatures that define the three resistance modes identified in the discrete simulations.

To further test whether introducing a mechanically distinct pericellular region could enable the FE model to reproduce observations in the discrete simulations, we manually defined a narrow pericellular shell (Fig. 8E) and varied its stiffness relative to the bulk matrix (*E*shell / *E*bulk) between 1 to 10. Increasing shell stiffness amplified local force magnitudes and produced partial localization of tensile forces near the poles. When shell stiffness was increased 10-fold, stresses became more strongly confined within the shell (Fig. 8G), reflecting reduced force transmission into the bulk matrix. Nevertheless, even at the highest stiffness ratio, the FE model did not reproduce the coexistence of radial compression and circumferential tension at the poles (Figs. 3A, B and 8G, H). Forces were predominantly tensile along the shell, and load-bearing paths, which were observed near the cell equator in discrete simulations, were absent. This discrepancy reflects an intrinsic limitation of continuum FEM: without microstructural degrees of freedom, it is hard to capture fiber alignment, buckling, or localized stress redistribution. Although shell stiffening increases force magnitude near the poles, continuum FE approaches are hard to reproduce the multimodal, coexisting force patterns that emerge naturally in discrete fibrous matrices.

## DISCUSSION

Physical interactions between dividing cells and their surrounding ECM continuously take place in biological processes, such as tissue remodeling and disease progression. Although the intracellular machinery that drives mitotic rounding and elongation has been characterized relatively well (*11, 12, 31, 32*), extracellular resistance has typically been reduced to bulk stiffness or simple geometric confinement in both experiments and models (*33–36*). Continuum-based approaches, which treat highly fibrous matrices as homogeneous elastic solids that oppose mitotic deformation uniformly, inherently assume that the ECM opposes mitotic elongation uniformly and in direct proportion to its stiffness. Yet, collagen-and fibrin-rich ECMs are anything but uniform: they are nonlinear, viscoelastic networks that stiffen under tension or shear, soften under compression, and relax forces over time. Cells actively exploit these microstructural nonlinearities during migration and spreading (*37–43*). However, it has remained largely unresolved how these same complex ECM properties resist mitotic shape changes.

In this study, we combined discrete, fiber-level modeling with live-cell imaging to define the microstructural origins of ECM resistance that cells encounter during cell growth and mitotic elongation (Figs. 1, 2). We found that the ECM does not behave as a static scaffold but instead acts as a mechanically adaptive material whose resistance is spatially patterned by cell shape changes and the mechanical history of the cell cycle (Figs. 3, 4). Prior to mitotic elongation, two-fold volumetric growth compacted and prestressed the proximal matrix, generating a densified pericellular shell that primarily resisted further volumetric expansion (Fig. 9A). During mitotic elongation, three distinct modes of ECM resistance emerged: (i) compressive resistance at the cell poles, (ii) resistance to areal expansion of the densified pericellular shell, and (iii) tensile resistance from equatorial fibers that couple the shell to the more distant matrix. Together, these modes function as a mechanical checkpoint on mitotic elongation and cell division. Because tension accelerates cross-linker dissociation, resistance from both the pericellular shell and its anchorage relaxes over time, whereas polar compressive resistance weakens through fiber buckling. Although all three resistance modes arose even without pre-mitotic volumetric growth, the pericellular shell contributed far less because it lacked prestress and densification (Fig. S1). These findings indicate that the ECM operates as a history-dependent mechanical checkpoint rather than a material with fixed, time-invariant properties.

**Figure 9.**
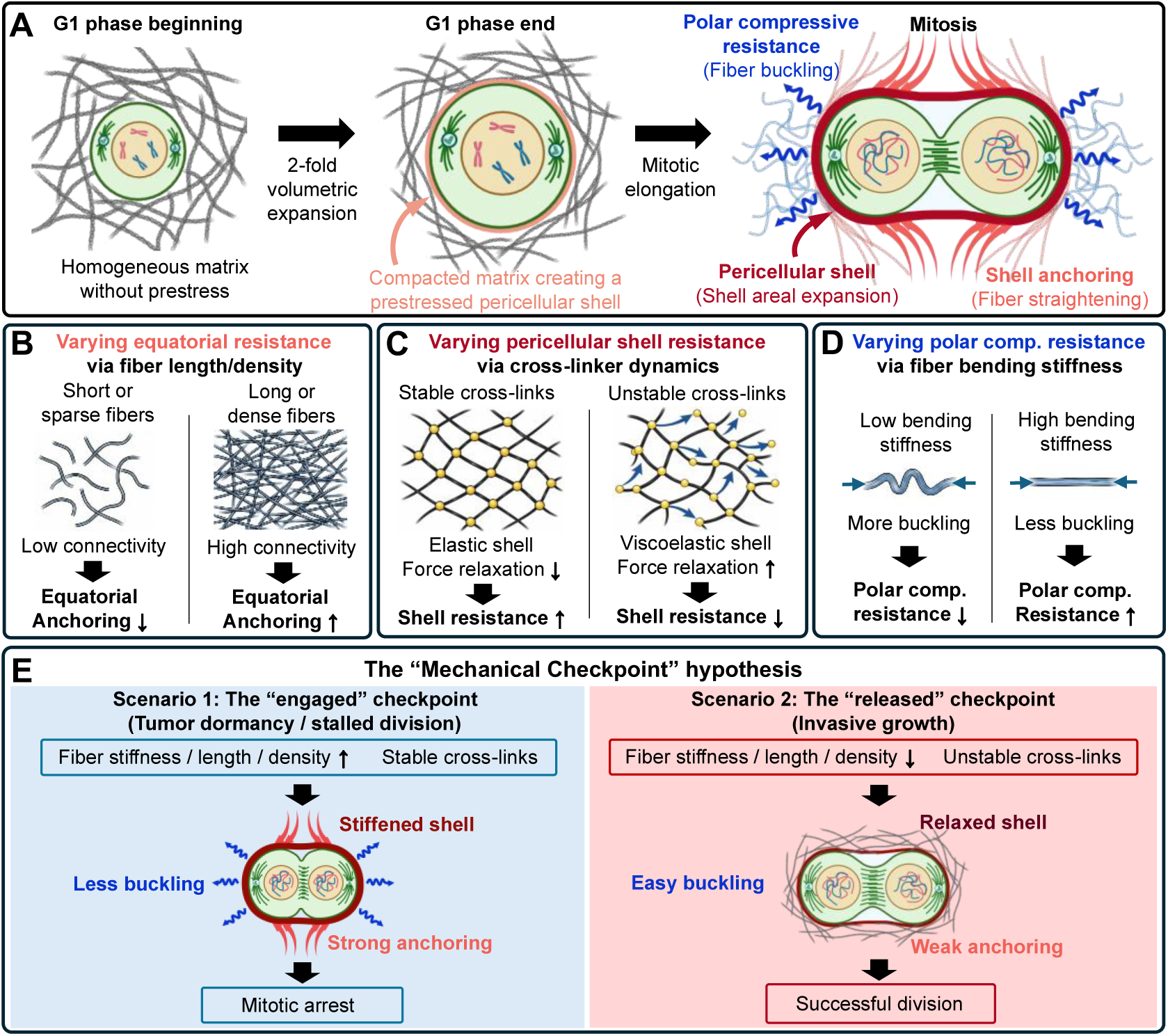
Growth-induced shell formation and extracellular matrix (ECM) microstructure set the mechanical resistance landscape for mitotic elongation. (A) Pre-mitotic volumetric growth during the G1 phase compacts an initially homogeneous ECM, generating a pre-stressed pericellular shell. Then, the ECM resists mitotic elongation in three distinct resistance modes: polar compressive resistance (blue), arising from fiber compression and buckling near the poles; pericellular shell resistance (red), originating from resistance to shell areal expansion; and equatorial anchoring (light red), arising from tensile loading and fiber straightening. (B) Equatorial anchoring is regulated by network connectivity, which is determined by fiber length and density: short or sparse fibers limit force transmission and weaken anchoring, whereas long or dense fibers enhance force transmission and strengthen anchoring. (C) Pericellular shell resistance is affected by cross-linker dynamics: stable cross-links relax shell tension less and thus increase resistance, whereas unstable cross-links promote force relaxation and reduce shell resistance. (D) Polar compressive resistance is regulated by fiber bending stiffness: compliant fibers buckle more readily and provide lower resistance, whereas stiffer fibers resist buckling and increase polar resistance. (E) The “Mechanical Checkpoint” hypothesis for tumor dormancy versus invasive growth. In the “engaged” checkpoint state, a matrix consisting of stiffer, longer, and denser fibers and stable cross-links significantly enhances all or some of the three resistance modes, potentially resulting in mitotic arrest and dormancy. In the “released” checkpoint state, a matrix composed of softer, shorter, and sparser fibers and unstable cross-links provides weaker mechanical resistance below the threshold required to impede mitotic elongation, enabling successful cell division and invasive growth.

Our analysis revealed that fiber density and length critically regulate the magnitude and distribution of mitotic resistance (Figs. 6 and S4A, B). With lower fiber density, cell elongation level was enhanced due to reduced matrix stiffness, which is similar to our previous experimental observations with different collagen concentrations (*44*), but the relative contribution of three types of resistances to mitotic elongation minimally changed. We also uncovered a percolation-like threshold affected by fiber length: when fibers were too short to form a continuous load-bearing network, the pericellular shell failed to connect effectively to the far-field matrix (Fig. 9B), rendering tensile anchorage negligible as in the case where the shell was intentionally disconnected from the far-field matrix (Fig. S1). Under such fragmented conditions with short fibers, cells may evade the mechanical checkpoint imposed by the ECM during division, thereby enhancing proliferative capacity. This finding also suggests that engineered biomaterials with tunable fiber length could be used to modulate mitotic resistance, offering a potential strategy to influence cell-cycle progression and proliferation in 3D culture systems (*45*).

We also found that cross-linking emerges as a key regulator of ECM resistance. A change in cross-linking density resulted in similar effects caused by a variation in fiber density. (Fig. S5); fewer cross-linking points between fibers increased cell elongation level but hardly changed the relative contribution of each resistance. The transient and force-dependent behavior of cross-linkers affected ECM resistance in a more interesting manner (Figs. 7A-D and S4C, D); with faster cross-linker turnover, cells could elongate more, and the contribution of shell anchorage became smaller. When cross-linkers exhibit rapid turnover, the matrix can dissipate and redistribute stress more efficiently. This mechanical fluidization reduces effective confinement around the dividing cell and facilitates elongation (Fig. 9C). In particular, the shell anchorage can be disrupted more easily since it is formed only by a few cross-linked fibers. Notably, this behavior parallels the unjamming transition observed in collective cancer migration, suggesting that dividing cells can actively fluidize their surroundings to bypass mechanical checkpoints without matrix degradation (*46–50*). By contrast, when cross-links are long-lived or mechanically stabilized under load, the matrix behaves more elastically and retains stored stress (*51–55*), thereby increasing resistance to elongation and delaying mitotic progression. In biopolymer matrices such as collagen and fibrin, mechanical loading can promote cross-link unbinding or slippage, leading to strain-softening or stress relaxation (*24, 56*), whereas enzymatic cross-linking reinforces the network, increasing stiffness and confinement (*57, 58*). Together, these findings highlight a dual role for cross-link dynamics, suggesting that cells do not passively respond to the ECM but actively regulate it through mechanochemical feedback. Under restrictive conditions, cells may remodel cross-links to create space for division (*12*), whereas in other contexts they may promote cross-link stabilization to reinforce tissue architecture (*59*).

In addition, we probed how fiber stiffness affects ECM confinement (Figs. 7E-H and S6). When fiber bending stiffness was increased 10-fold, cell elongation was reduced to highly increased compressive resistance at poles (Fig. 9D). The fiber bending stiffness is roughly proportional to the fourth power of fiber thickness if fibers are treated as a beam. In case of collagen, fiber thickness can vary over a wide range, depending on conditions (*60*). With the same collagen concentration, if fiber thickness increases, fiber density decreases as [thickness]^-2^. Then, a decrease in ECM resistance due to lower overall matrix stiffness can be compensated by the enhanced compressive resistance at poles, meaning the major mode of ECM resistance is switched. In addition, a change in fiber extensional stiffness had a noticeable effect on ECM resistance. When fibers were highly stretchable, shell anchorage played a less significant role because the anchorage relies on a few fibers unlike the shell consisting of more fibers; the effective extensional stiffness of the shell would be much higher than that of a single fiber. Elastin fibers are known to be much softer than collagen fibers (i.e., lower Young’s modulus) (*61*), so their bending and extensional stiffnesses are much lower. Thus, it is expected that a matrix with a high fraction of elastin resists mitotic elongation primarily via the pericellular shell. From a biomaterial perspective, these findings highlight opportunities to engineer mechanoadaptive scaffolds that dynamically regulate mechanical confinement throughout the cell division cycle (Fig. 9D). Incorporating stimuli-responsive or reversible cross-linkers could enable tunable transitions between stiffening and softening, effectively engaging or releasing the mechanical checkpoint on demand—restricting proliferation in tumor-like microenvironments or permitting mitosis during regeneration within dense tissues (*62*). Guided by our observation that long-range force transmission is a prerequisite for ECM resistance, we propose that regenerative scaffolds should adopt fiber architectures below the percolation threshold, thereby minimizing mechanical braking and accelerating proliferation. Conversely, tumor-mimetic models must maintain sufficient connectivity to enforce dormancy. Importantly, our results show that ECM resistance is shaped not only by forces generated during mitotic elongation but also by pre-mitotic volumetric growth, which densifies and prestresses the surrounding matrix. Scaffolds that constrain swelling may intensify confinement and hinder elongation, whereas materials that permit controlled expansion can reduce resistance and support successful division. Thus, porosity, swelling behavior, elasticity, and cross-linking dynamics must be co-optimized to accommodate the full temporal sequence of cell division and to enable precise control over proliferation in 3D culture, regenerative systems, and disease-mimetic models.

We confirmed that the unique resistance modes, which we found, arise from the polymeric nature of fibrillar matrices by comparing with finite-element simulations based on hyperelastic constitutive laws (Fig. 8). Although the continuum models reproduced the overall deformation fields around a dividing cell, they failed to capture two key features: the emergence and mechanical resistance of the pericellular shell, and the tensile anchorage between this shell and the distant matrix. This limitation stems from the fact that continuum approaches are hard to represent the non-affine, topology-driven fiber rearrangements that occur in sparse, disordered networks. This comparison between the two modeling frameworks also underscores that discrete fiber-level representations are better for predicting how the surrounding matrix responds to mitotic forces.

Admittedly, our study has some limitations. Our extensive parametric analysis relied on 2D approximations to maintain computational feasibility although we confirmed the universality of the mechanical checkpoint using a fully 3D fiber-resolution model (Figs. 5 and S3). Extending these high-throughput sweeps into fully 3D geometries will be essential for capturing emergent topological features—such as fiber entanglement, out-of-plane buckling, and isotropic percolation—that may further amplify confinement in vivo. Additionally, we excluded enzymatic degradation in our model to isolate the baseline mechanical barrier. Coupling this framework with proteolytic kinetics would enable direct quantitative comparisons between viscous slippage and matrix degradation as competing routes for overcoming confinement during cell division. Lastly, our model does not account for any signaling pathways that may play a role during cell division. Integrating mechanotransductive feedback loops, such as YAP/TAZ signaling, could reveal how the high compressive stress of the pericellular shell drives cytoplasmic sequestration of YAP and promotes dormancy (*63–65*). Such extensions would bridge matrix-level mechanics with intracellular decision-making, providing a more complete picture of how cells navigate and potentially escape the mechanical constraints imposed by dense 3D microenvironments.

## CONCLUSIONS

Taken together, our study redefines the ECM not as a static barrier, but as a tunable mechanical checkpoint regulated by growth history and network topology. By shifting the focus from bulk stiffness to fiber architecture and cross-link dynamics, we provide a unified framework for understanding how cells negotiate physical confinement. These insights suggest that targeting the viscous properties of the ECM rather than just its stiffness could offer new therapeutic avenues to re-engage the mechanical checkpoint and induce dormancy in metastatic tumors.

## Notes

### Competing Interest Statement

The authors have declared no competing interest.

### Funding

- National Institute of General Medical Sciences: R01GM151628

