## Supplementary Materials for "Mechanical Checkpoint for Cell Division in Three-Dimensional Microenvironments"

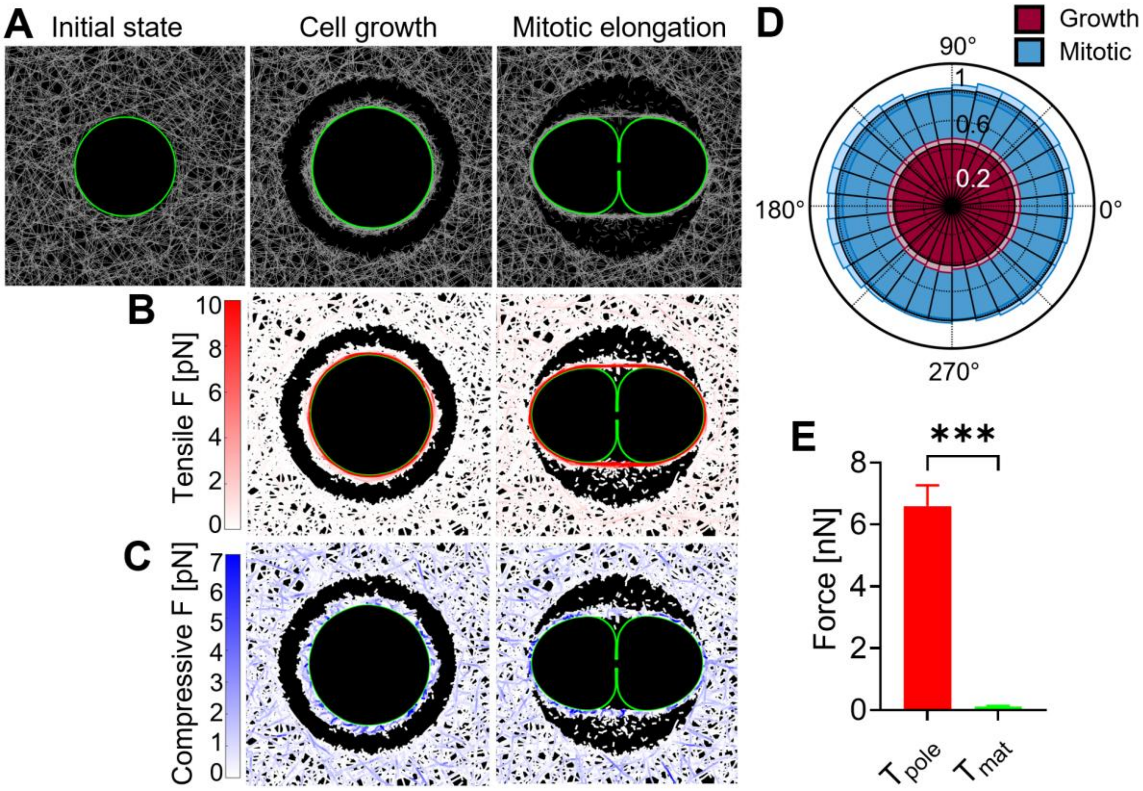


**Figure S1. A pericellular shell separated from a bulk matrix still strongly resisted mitotic elongation with isotropic tension development.** (A) Snapshots illustrating three phases: initial pre-growth, post-growth, and mitotic elongation. A 3-μm gap is created between the shell and the bulk matrix to isolate local mechanical interactions between the shell and the cell, with pericellular shell remaining intact, (B) Tensile forces acting on the matrix. In both cases, tensile forces were locally distributed along the shell. (C) Compressive forces acting on the matrix, showing no noticeable development. (D) Angular distribution of tensile forces along the shell at the end of the growth (red) and during mitotic elongation (blue). Lighter colors indicate standard deviations. (E) Quantitative comparison of tensile resistance components measured from the shell during mitotic elongation. $T_{pole}$ denotes the tensile force measured on the shell near the poles, and $T_{mat}$ is equal to a difference between $T_{pole}$ and the tensile force measured on the shell near the cell equator. Data in (D, E) are presented as mean ± s.d. ($n$ = 4 independent simulations). Statistical analysis in (E) was performed using two-sided unpaired t-tests (C). ***: $p < 0.001$.

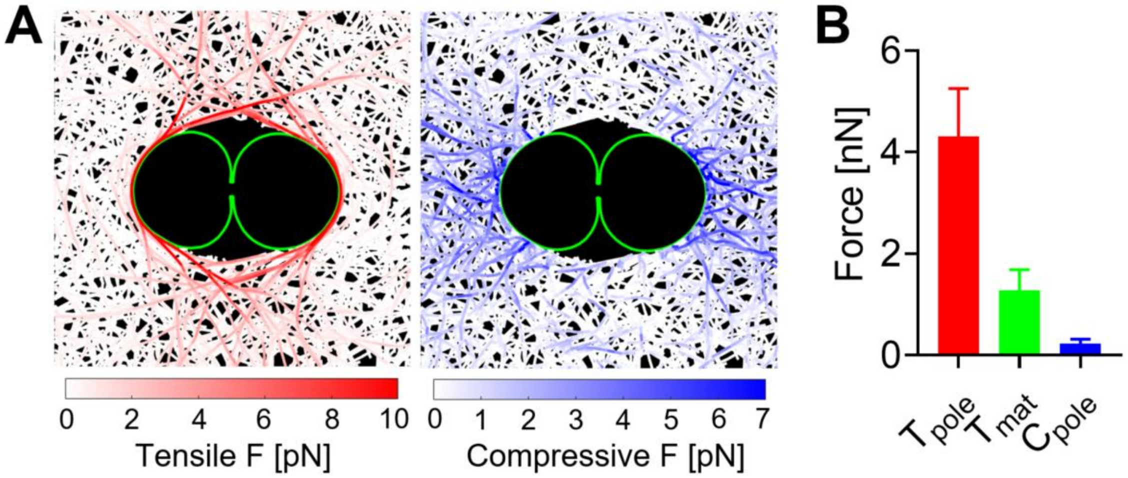


**Figure S2. Mitotic elongation without pre-mitotic growth resulted in the three resistance modes but to a lesser extent.** (A) Snapshot showing tensile (left) and compressive (right) forces acting on a matrix. The absence of volumetric cell expansion before mitotic elongation still led to similar force development patterns but resulted in enhanced cell elongation, larger matrix voids, and weaker anchor development. (B) Quantification of three resistance modes: $T_{\text{pole}}$ indicating the tensile force measured on the shell near the poles, $T_{\text{mat}}$ representing the relative contribution of anchoring between the shell and a bulk matrix, and $C_{\text{pole}}$ indicating compressive forces near the poles.

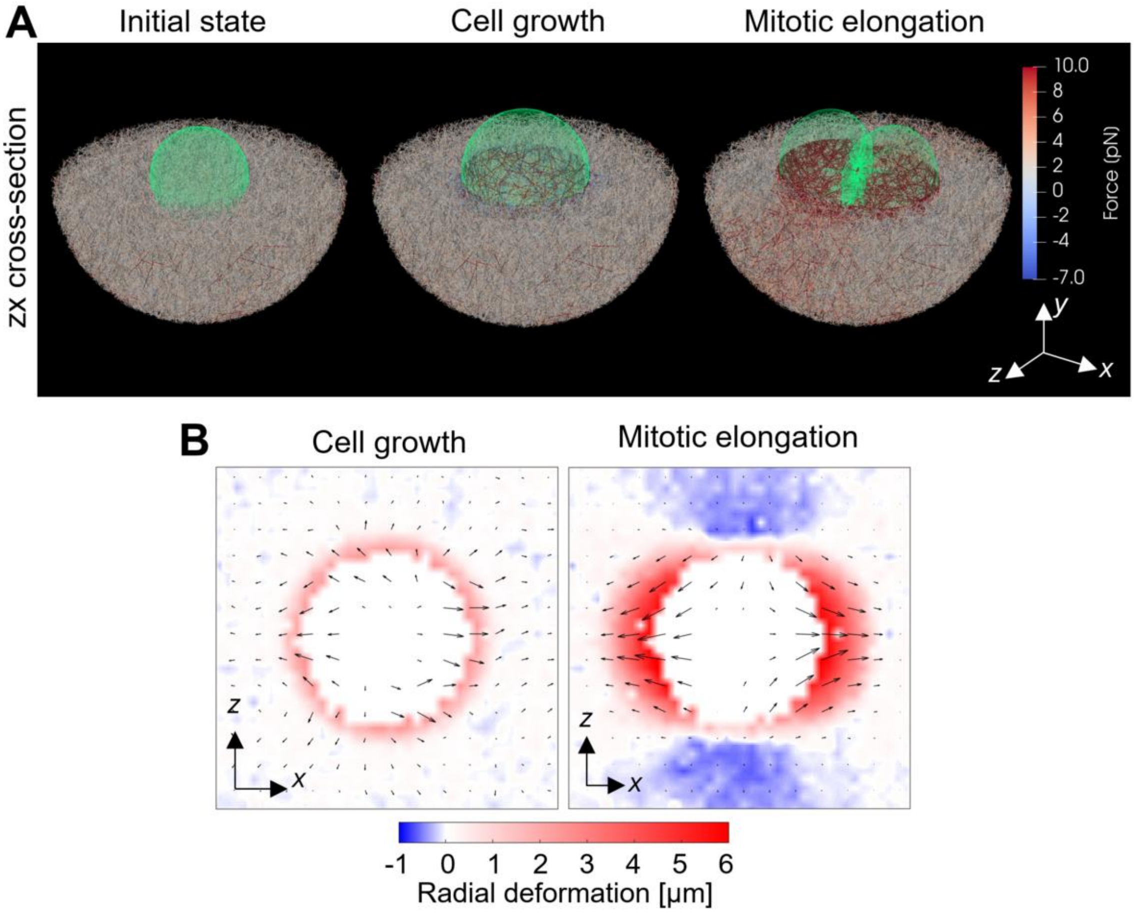


**Figure S3. Additional data from the three-dimensional simulations.** The same simulation data were used as those used in Fig. 5. (A) Snapshots showing the three stages of cell division in 3D simulations: pre-growth, post-growth, and mitotic elongation, using their zx cross-sections. The cell boundary is shown in green, and color scaling on the right is used to visualize forces acting on matrix fibers. (B) Matrix deformation fields at the end of the growth phase and during mitotic elongation, measured on the zx cross-sections.

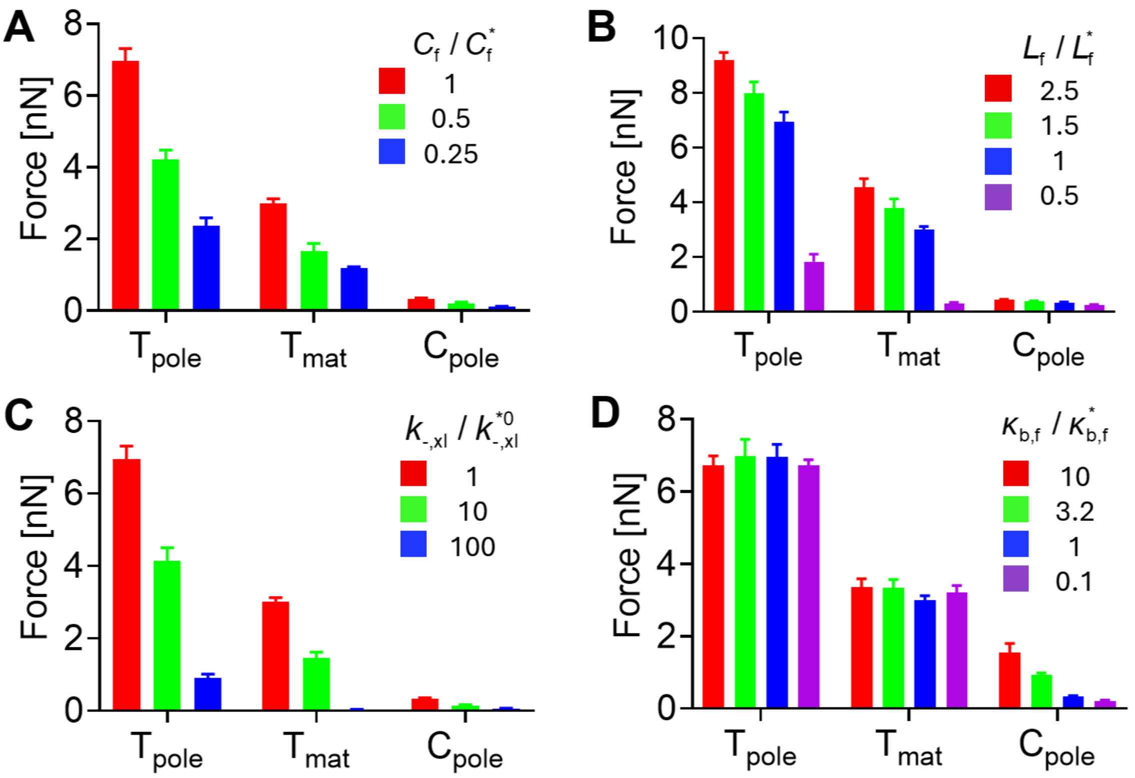


**Figure S4. Raw data of forces representing the three resistance modes with a change in matrix parameters:** (A) fiber density ($C_f$), (B) fiber length ($L_f$), (C) the unbinding rate of cross-linkers ($k^0_{-,xl}$), and (D) fiber bending stiffness ($\kappa_{b,f}$). The subscript "*" denotes their reference values. The same simulation data were used as those used in Figs. 6 and 7.

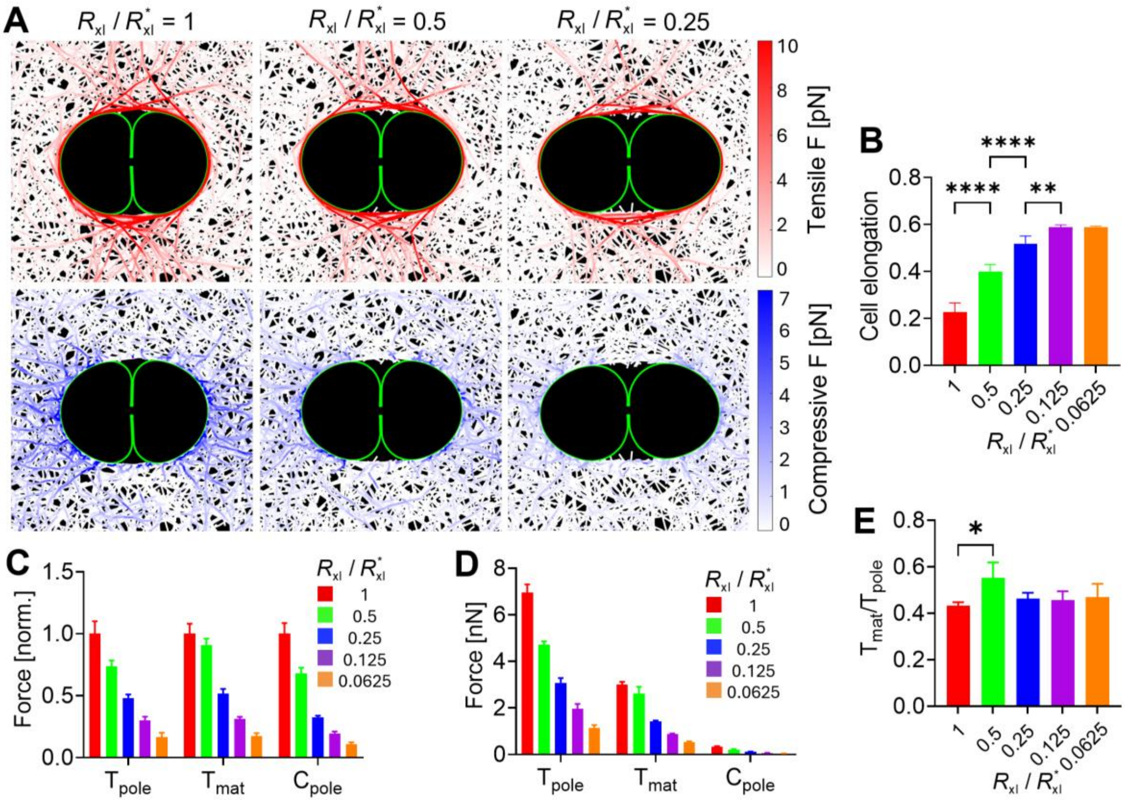


**Figure S5. Reduced cross-linking level between fibers decreased matrix resistance and thus enhanced mitotic cell elongation.** Cross-linking density ($R_{xl}$) was reduced from its reference value ( $R^*_{xl}$ ). (A) Snapshots showing the spatial distribution of tensile (top) and compressive (bottom) forces acting on a matrix with varying $R_{xl}$. (B) Cell elongation as a function of $R_{xl}$. Lower $R_{xl}$ led to greater cell elongation. (C, D) Analysis of forces representing three resistance modes, depending on $R_{xl}$: shell tensile resistance ($T_{pole}$), anchor resistance ($T_{mat}$), and polar compressive resistance ($C_{pole}$). (E) The ratio of $T_{mat}$ to $T_{pole}$ which represents the relative importance of anchors between the pericellular shell and the bulk matrix. Overall, the three types of forces decreased to similar extents in proportion to a decrease in $R_{xl}$, meaning the relative contribution of those three resistance modes did not change much by the variation in $R_{xl}$. Interestingly, halving $R_{xl}$ increased the relatively importance of the anchors by ~25%.

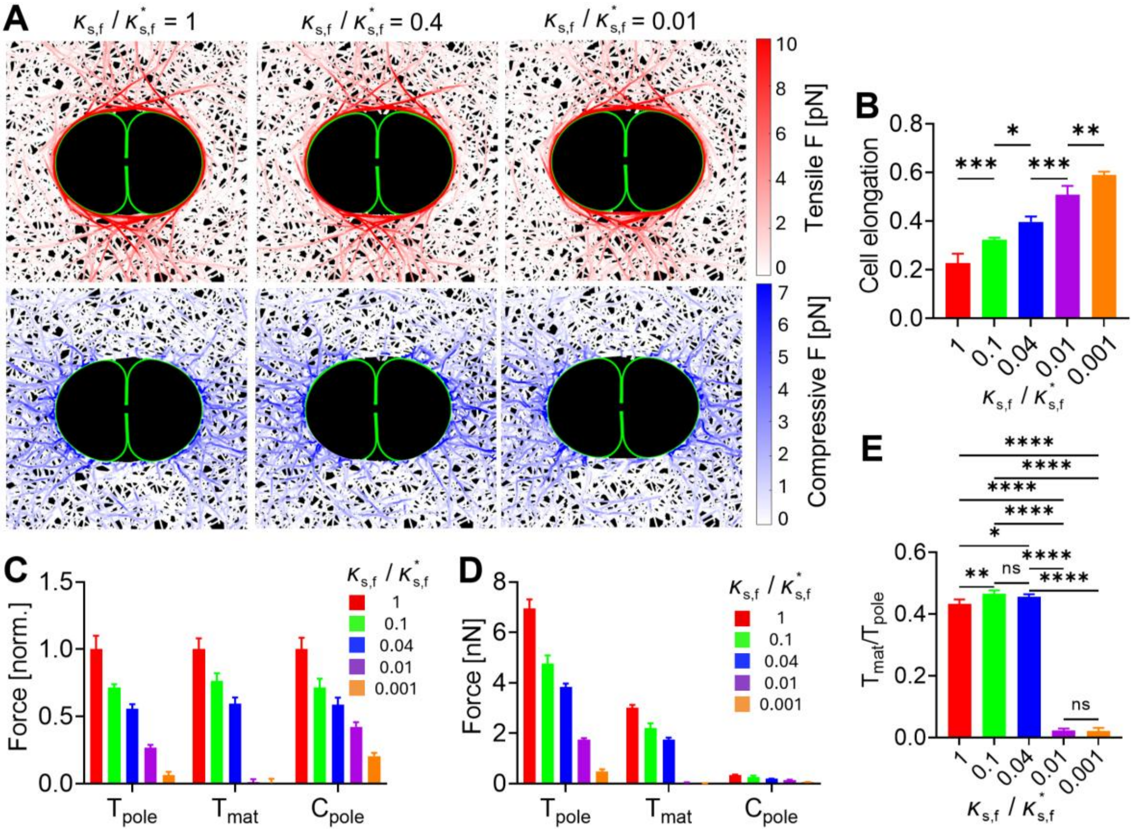


**Figure S6. Lower extensional stiffness of fibers ($\kappa_{s,f}$) enhanced mitotic elongation by reducing matrix resistance, in particular, anchor resistance.** $\kappa_{s,f}$ was reduced from its reference value ($\kappa^{*}_{s,f}$). (A) Snapshots visualizing the spatial distribution of tensile (top) and compressive (bottom) forces exerted on a matrix with different $\kappa_{s,f}$. (B) Cell elongation, depending on $\kappa_{s,f}$. Lower $\kappa_{s,f}$ resulted in higher cell elongation. (C, D) Quantification of forces representing three types of matrix resistance with various $R_{xl}$: shell tensile resistance ($T_{pole}$), anchor resistance ($T_{mat}$), and polar compressive resistance ($C_{pole}$). (E) The ratio of $T_{mat}$ to $T_{pole}$ which indicates the relative significance of anchors between the pericellular shell and the bulk matrix. When $\kappa_{s,f}$ was reduced up to 1/25 of $\kappa^{*}_{s,f}$, all three resistance forces were decreased to similar extents. However, when $\kappa_{s,f}$ decreased further, $T_{mat}$ became negligible compared to $T_{pole}$, implying that the shell was no longer mechanically anchored to the matrix. Interestingly, $C_{pole}$ showed dependence on $\kappa_{s,f}$ although fiber buckling is affected mainly by fiber bending stiffness ($\kappa_{b,f}$).

**Supplementary Table 1.** List of parameters employed in the discrete fiber-level simulations. The subscript "*" indicates the reference value of each parameter.

| Symbol | Definition | Value |
|---|---|---|
| $r_{0,\mathrm{f}}$ | Length of fiber segments | $1.0\times10^{-6}$ [m] |
| $r_{\mathrm{c,f}}$ | Diameter of fibers | $2.0\times10^{-7}$ [m] |
| $\theta_{0,\mathrm{f}}$ | Bending angle formed by serially connected fiber segments | 0 [rad] |
| $\kappa_{\mathrm{s,f}}$ | Extensional stiffness of fibers | $4.0\times10^{-3}$ [N/m]* |
| $\kappa_{\mathrm{b,f}}$ | Bending stiffness of fibers | $4.14\times10^{-19}$ [N·m]* |
| $r_{0,\mathrm{xl}}$ | Length of cross-linker arms | $2.0\times10^{-7}$ [m] |
| $r_{\mathrm{c,xl}}$ | Diameter of cross-linker arms | $2.0\times10^{-8}$ [m] |
| $\theta_{0,\mathrm{xl},1}$ | Bending angle formed by two cross-linker arms | 0 [rad] |
| $\theta_{0,\mathrm{xl},2}$ | Bending angle formed by a cross-linker arm and the axis of a fiber where the arm is bound | π/2 [rad] |
| $\kappa_{\mathrm{s,xl}}$ | Extensional stiffness of cross-linkers | $2.0\times10^{-3}$ [N/m] |
| $\kappa_{\mathrm{b,xl}}$ | Bending stiffness of cross-linkers | $1.04\times10^{-19}$ [N·m] |
| $r_{0,\mathrm{m}}$ | Initial size of membrane elements | $4.0\times10^{-7}$ [m] |
| $r_{\mathrm{c,m}}$ | Thickness of membrane elements | $3.0\times10^{-7}$ [m] |
| $\theta_{0,\mathrm{m}}$ | Bending angle formed by adjacent membrane elements | 0 [rad] |
| $\kappa_{\mathrm{s,m}}$ | Extensional stiffness of the membrane | $1.0\times10^{-4}$ [N/m] |
| $\kappa_{\mathrm{b,m}}$ | Bending stiffness of the membrane | $1.0\times10^{-18}$ [N·m] |
| $\kappa_{\mathrm{r,m}}$ | Strength of repulsive force | $4.0\times10^{-4}$ [N/m] |
| $\kappa_{\mathrm{v,m}}$ | Strength of volume conservation | 100 [Pa] |
| $C_{\mathrm{f}}$ | Fiber concentration | 1 [μM] |
| $L_{\mathrm{f}}$ | Average length of fibers | ~10 [μm]* |
| $R_{\mathrm{xl}}$ | Ratio of # of cross-linkers to # of fibers | ~2.86 [cross-linker/fiber]* |
| $N_{\mathrm{m}}$ | Number of membrane nodes | 158 (2D) / 10,242 (3D) |
| $\Delta t$ | Time step | $3.97\times10^{-4}$ [s] |
| $\mu$ | Viscosity of a medium | 8.6 [Pa·s] |
| $k_{+,\mathrm{xl}}$ | Binding rate constant of cross-linkers | 0.1 [$s^{-1}$] |
| $k^{0}_{-,\mathrm{xl}}$ | Zero-force unbinding rate constant of cross-linkers | $1.0\times10^{-6}$ [$s^{-1}$] |
| $\lambda_{-,\mathrm{xl}}$ | Force sensitivity of cross-linker unbinding | $4.0\times10^{-10}$ [m] |
| $k_{\mathrm{B}}T$ | Thermal energy | $4.14\times10^{-21}$ [J] |
| $F_{\mathrm{CRC}}$ | Force representing the cytokinetic ring constriction | $2.0\times10^{-10}$ [N] |
| $F_{\mathrm{ISE}}$ | Force representing the interpolar spindle elongation | $1.0\times10^{-10}$ [N] |

**Supplementary Table 2.** List of parameters used in finite element simulations.

| Symbol | Definition | Value |
|---|---|---|
| $R_{\mathrm{cell}}$ | Radius of the cell | $1.0\times10^{-5}$ [m] |
| $E_{\mathrm{cell}}$ | Initial elastic modulus of the cell | 324 [$N/m^2$] |
| $\nu_{\mathrm{cell}}$ | Poisson's ratio of the cell | 0.5 (incompressible) |
| $R_{\mathrm{shell}}$ | Radius of the pericellular shell | $1.05\times10^{-5}$ [m] |
| $E_{\mathrm{shell}}$ | Initial elastic modulus of the pericellular shell | 8320 [$N/m^2$] |
| $\nu_{\mathrm{shell}}$ | Poisson's ratio of the pericellular shell | 0.313 |
| $R_{\mathrm{mat}}$ | Radius of the matrix | $5.0\times10^{-5}$ [m] |
| $E_{\mathrm{mat}}$ | Initial elastic modulus of the matrix | 832 [$N/m^2$] |
| $\nu_{\mathrm{mat}}$ | Poisson's ratio of the matrix | 0.313 |
